# G protein-coupled potassium channels implicated in mouse and cellular models of GNB1 Encephalopathy

**DOI:** 10.1101/697235

**Authors:** Sophie Colombo, Sabrina Petri, Boris Shalomov, Haritha P. Reddy, Galit Tabak, Ryan S. Dhindsa, Sahar Gelfman, Sasa Teng, Daniel Krizay, Elizabeth E. Rafikian, Amal K. Bera, Mu Yang, Michael J. Boland, Yueqing Peng, Wayne N. Frankel, Nathan Dascal, David B. Goldstein

## Abstract

*De novo* mutations in *GNB1*, encoding the Gβ1 subunit of G proteins, cause a neurodevelopmental disorder with global developmental delay and epilepsy. Mice carrying a pathogenic mutation, K78R, recapitulate aspects of the disorder, including developmental delay and frequent spike-wave discharges (SWD). Cultured mutant cortical neurons display aberrant bursting activity on multi-electrode arrays. Strikingly, the antiepileptic drug ethosuximide (ETX) restores normal neuronal network behavior *in vitro* and suppresses SWD *in vivo*. In contrast, while valproic acid suppresses SWD, it does not restore normal network behavior, suggesting that ETX has mechanistic specificity for the effects of aberrant Gβ1 signaling. Consistent with this, we show that K78R is a gain-of-function of G protein-coupled potassium channel (GIRK) activation that is potently inhibited by ETX. This work suggests that altered Gβ1 signaling causes disease in part through effects on GIRK channels, illustrates the utility of cultured neuronal networks in pharmacological screening, and establishes effective pre-clinical models for GNB1 Encephalopathy.

## INTRODUCTION

G protein-coupled receptors (GPCRs) constitute the largest family of transmembrane receptors, regulating many aspects of human physiology (Rosenbaum et al., 2009). Most GPCR signaling is transduced through a G protein heterotrimeric complex, composed of three subunits, Gα, Gβ and Gγ. GPCR activation catalyzes a GDP to GTP exchange on the Gα subunit, leading to the dissociation of Gα from the obligate dimer Gβγ (Oldham and Hamm, 2006, 2008; Smrcka, 2008). Both Gα and Gβγ then bind and regulate a wide variety of downstream effectors (Khan et al., 2013; Khan et al., 2016; Smrcka, 2008).

Mutations in *GNB1*, encoding the Gβ1 subunit of G proteins, cause GNB1 Encephalopathy, a severe neurodevelopmental disorder characterized by global developmental delay, speech and ambulatory deficits, intellectual disability and a variety of seizure types (Hemati et al., 2018; Petrovski et al., 2016).

Unfortunately, the signaling pathways mediated by Gβγ are broad and complex, presenting a challenge to hypothesizing *GNB1* disease mechanims. Indeed, neurological functions of Gβγ dimers include regulation of voltage-dependent calcium Ca_v_ channels (VDCCs) (Dascal, 2001; Dolphin, 1998; Ikeda, 1996; Weiss, 2010; Zamponi and Currie, 2013), postsynaptic G protein-coupled inwardly-rectifying potassium channels (GIRK) (Chen and Johnston, 2005; Luscher and Slesinger, 2010; Malik and Johnston, 2017), and vesicle release through interaction with the SNARE complex (Alford et al., 2018; Yoon et al., 2007; Zurawski et al., 2017; Zurawski et al., 2016).

To elucidate how *GNB1* mutations cause disease, we generated a mouse model of the K78R human pathogenic variant using CRISPR/Cas9. *Gnb1*^K78R/+^ mice recapitulate many clinical features of affected individuals, including developmental delay, motor and cognitive deficits, and absence-like generalized seizures. Cortical neurons from *Gnb1*^K78R/+^ mice cultured *in vitro* also display extended bursts of firing followed by extensive recovery periods. Both the *in vivo* and *in vitro* excitability phenotypes are effectively suppressed by the anti-epileptic drug ethosuximide (ETX). Finally, we expressed wild-type and mutant Gβ1 with GIRK1/2 and GIRK2 channels in *Xenopus* oocytes and show that the K78R mutation increases GIRK activation, and that ETX has its effects in part through GIRK channel inhibition.

## RESULTS

### Characterization of the *Gnb1*^K78R/+^ mouse model and Gβ1 expression

We modeled the first mutation identified in a patient, K78R, using CRISPR/Cas9. The K78R knock-in (KI) mutation (human NM_002074, mouse NM_008142, c.233 A>G; p.K78R) was verified by Sanger sequencing and PCR (Figure S1A-C). We also obtained a knock-out (KO) line with a 71bp deletion (Figure S1D). Both KO and KI lines showed homozygous embryonic lethality (Figure S1E). However, while heterozygous KO pups are viable, heterozygous KI pups, *Gnb1*^K78R/+^, showed reduced viability on an isogenic C57BL/6NJ (B6NJ) background, and a F1 Hybrid (F1H) background (FVB.129P2 x C57BL/6NJ) (Figure S1E). We observed normal Gβ1 protein levels in postnatal day 0 (P0) *Gnb1*^K78R/+^ cortex compared to WT, with high Gβ1 expression at the membrane and low expression in the cytosol (Figure S1F,G). Taken together, these observations demonstrate that the K78R mutation does not confer a null allele and that the protein is properly targeted to the membrane.

We also assessed Gβ1 expression in primary cortical neurons from WT and *Gnb1*^K78R/+^ pups. Gβ1 is expressed in both excitatory and inhibitory neurons (Figure S2A-C), but is not expressed in astrocytes (Figure S2D). Gβ1 preferentially localizes in the soma, near the plasma membrane, and is not present in the axon initial segment (Figure S2E,E’). Gβ1 is expressed in both deep and upper layer cortical neurons (Figure S2F). We observed similar Gβ1 expression levels and localization patterns in WT and *Gnb1*^K78R/+^ neurons (Figure S2E’ and data not shown).

### *Gnb1*^K78R/+^ mice exhibit phenotypes relevant to clinical features of GNB1 Encephalopathy

Both male and female *Gnb1*^K78R/+^ pups had a significantly lower body weight than WT littermates at birth (Figure 1A,B). To assess behavioral phenotypes pertinent to key clinical features of GNB1 Encephalopathy, we tested *Gnb1*^K78R/+^ pups for physical and sensorimotor developmental milestones between postnatal days (P) 4 and 11, and *Gnb1*^K78R/+^ adult mice for motor and cognitive functions. Early developmental milestones including body weight, surface righting reflex, negative geotaxis, vertical screen holding, and separation-induced ultrasonic vocalizations (USV) were assayed as previously described (Yang et al., 2012). On a B6NJ background, *Gnb1*^K78R/+^ pups weighed significantly less than WT littermates across early postnatal days (Figure 1C), and remained significantly smaller as young adults at 6 weeks (Figure S3A). *Gnb1*^K78R/+^ pups exhibited deficits in the surface righting reflex test, being initially faster than WT littermates at P4 but not progressing afterwards, and had a significant delay in both 90⁰ and 180⁰ negative geotaxis tests. They did not however show deficits in vertical screen holding, suggesting normal grip strength (Figure 1C). Finally, while WT pups presented a classic inverted U-shaped curve of separation-induced USV, *Gnb1*^K78R/+^ pups emitted few vocalizations throughout the tested days, indicating abnormal development (Figure 1C). Pups on a F1H background are generally bigger and healthier than on a B6NJ background. *Gnb1*^K78R/+^ pups presented similar, albeit milder, developmental delays compared to WT littermates in this background (Figure S3B).

**Figure 1.**
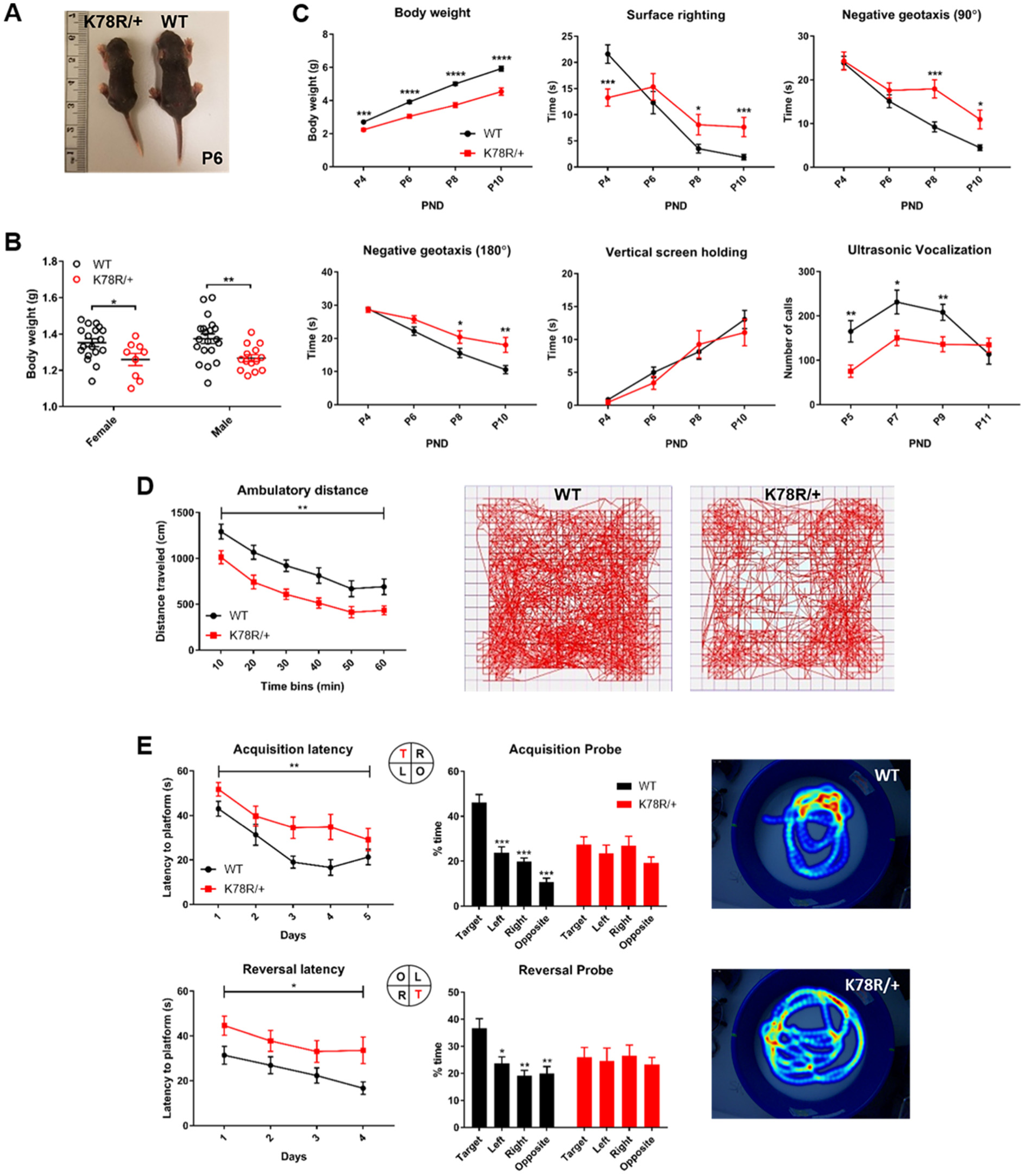
*Gnb1*^K78R/+^ mice exhibit developmental delay as neonates, and motor and cognitive deficits as adults. **(A)** A representative photo of a *Gnb1*^K78R/+^ pup (left) and a WT littermate (right) at P6 on the B6NJ background. **(B)** Pup weight at P0 on the B6NJ background. WT (black circles): n = 18 females and 20 males; *Gnb1*^K78R/+^ (red circles): n = 9 females and 14 males; females *p < 0.05 vs. WT and males **p < 0.01 vs. WT. Mann-Whitney U test with 10000 permutations. **(C)** Developmental milestones between P4 and P10 on the B6NJ background include body weight, surface righting reflex, negative geotaxis 90° and 180° from downward-facing position, and vertical screen holding: n = 32 WT and 20 *Gnb1*^K78R/+^. Separation-induced USV between P5 and P11: n= 28 WT and 20 *Gnb1*^K78R/+^. *p < 0.05; **p < 0.01; ***p < 0.001; ****p < 0.0001. Mann-Whitney U test with 10000 permutations. **(E)** Open field test on the B6NJ background. Distance traveled: n= 18 WT and 13 *Gnb1*^K78R/+^; **p < 0.01. Two-way repeated measures ANOVA. Representative movement traces of a WT (left) and a *Gnb1*^K78R/+^ (right) mouse walking for 1 hour. **(F)** Morris water maze test on the B6NJ background. n = 15 WT and 14 *Gnb1*^K78R/+^. Learning: **p < 0.01 for acquisition latency; *p < 0.05 for reversal latency. Two-way repeated measures ANOVA. Memory: ***p < 0.001 for Target vs. Left, Target vs. Right and Target vs. Opposite during acquisition probe trial; *p < 0.05 for Target vs. Left, **p < 0.01 for Target vs. Right and Target vs. Opposition during reversal probe trial. One-way repeated measures ANOVA with Bonferroni multiple comparisons test. Representative heat maps of a WT (top) and a *Gnb1*^K78R/+^ mouse during reversal probe trial. *All gra*p*hs represent mean ± SEM. PND: postnatal day.* See also Figure S3.

We assessed spontaneous locomotor activity in the open field, and observed that adult *Gnb1*^K78R/+^ mice were significantly less active compared to WT littermates (Figure 1D). Vertical exploration (rearing) and center time were not different (Figure S3C), suggesting that reduced activity is not attributable to changes in anxiety-like behaviors. *Gnb1*^K78R/+^ mice did not show deficits in the rotarod test, suggesting normal balance and grip strength (Figure S3D).

We tested adult mice for cognitive functions. While *Gnb1*^K78R/+^ mice did not exhibit deficits in the classic contextual and cued fear conditioning tests (Figure S3E), they displayed spatial learning and memory deficits in the Morris Water Maze test. These data could indicate that while *Gnb1*^K78R/+^ mice can recognize contextual cues, they are impaired in using spatial cues to navigate in a complex environment. In the water maze, *Gnb1*^K78R/+^ mice were slower at acquiring the spatial learning task and the subsequent reversal task (Figure 2E). Importantly, these deficits were observed with normal swimming speed in *Gnb1*^K78R/+^ mice (Figure S3F), indicating swimming ability did not confound learning performance in this test. *Gnb1*^K78R/+^ mice also displayed memory deficits. During the probe test of both acquisition and reversal, WT mice spent significantly more time in the trained quadrant than in the other three quadrants, whereas *Gnb1*^K78R/+^ mice did not show preference for the quadrant where the platform used to be (Figure 2E and Figure S3G). In conclusion, *Gnb1*^K78R/+^ mice exhibited behavioral phenotypes relevant to GNB1 Encephalopathy which is characterized by global developmental delay, ambulatory deficits and intellectual disability.

**Figure 2.**
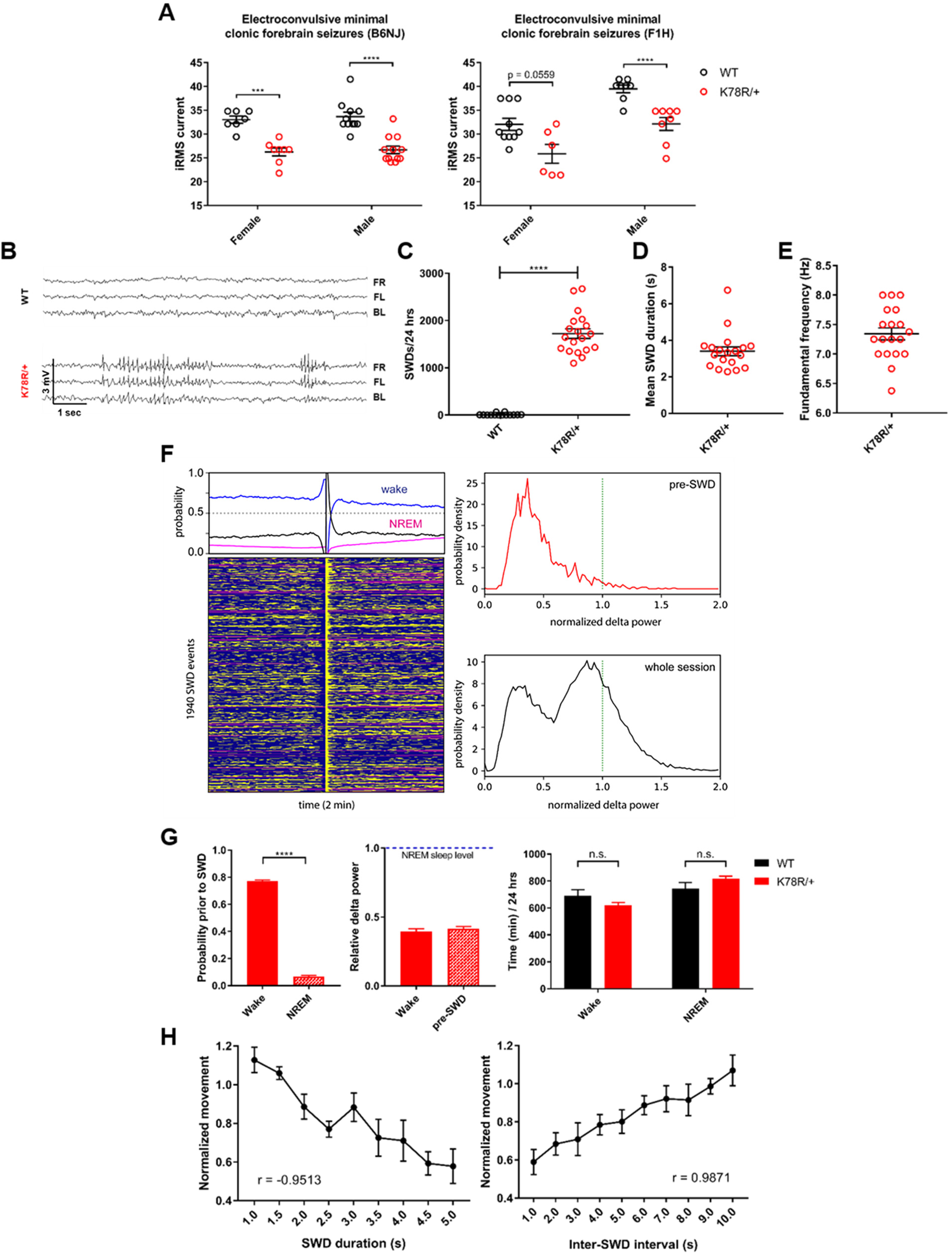
Excitability phenotypes in *Gnb1*^K78R/+^ mice. **(A)** Electroconvulsive threshold tests. Minimal clonic forebrain seizure endpoint in B6NJ (left) and F1H (right) mice. n (B6NJ) = 7 WT female, 11 WT male, 8 *Gnb1*^K78R/+^ female and 12 *Gnb1*^K78R/+^ male; n (F1H) = 10 WT female, 8 WT male, 6 *Gnb1*^K78R/+^ female and 8 *Gnb1*^K78R/+^ male; ***p < 0.001; ****p < 0.0001 Mann-Whitney U test with 10000 permutations. **(B)** Representative traces from video-EEG recordings of cortically implanted *Gnb1*^K78R/+^ and WT littermates on the F1H background showing SWDs in *Gnb1*^K78R/+^ mice. FR: front right, FL: front left; BL: back left. **(C)** Quantification of SWDs per 24 h. n = 14 WT and 19 *Gnb1*^K78R/+^; WT = 11.71 ± 6.527 vs. *Gnb1*^K78R/+^ = 1724 ± 100.6; ****p < 0.0001; Mann-Whitney U test with 10000 permutations. **(D)** Mean duration of SWDs. n = 19 *Gnb1*^K78R/+^; mean ± SEM: 3.397 ± 0.239. **(E)** Peak spectral power. n = 19 *Gnb1*^K78R/+^; mean ± SEM: 7.342 ± 0.101. **(F)** Correlation between SWD and sleep/wake states in a *Gnb1*^K78R/+^ mouse. Left, brain states 1 min before and 1 min after aligned 1940 SWD events that occurred over a 24 h period. Probabilities of each brain state are plotted on the top (NREM: purple; wake: blue; SWD: yellow). Right, probability density of normalized delta power in pre-SWD windows (3 s period prior to SWD onset, top) and in the whole 24 h session (bottom). Note that the second peak with larger delta power (indicator of NREM sleep) was absent in the pre-SWD window. **(G)** Probability of wake and NREM state prior to SWD. n = 19 *Gnb1*^K78R/+^; wake = 0.7371 ± 0.01 vs. NREM = 0.075 ± 0.007; ****p < 0.0001. Delta power during wake and pre-SWD, relative to NREM sleep. n = 19 *Gnb1*^K78R/+^; wake = 0.40 ± 0.02 vs. pre-SWD = 0.42 ± 0.015; n.s.. Time spent in wake or NREM over a 24 h period. n = 12 WT and 19 *Gnb1*^K78R/+^; WT vs. *Gnb1*^K78R/+^: n.s.; Mann-Whitney U test with 10000 permutations. **(H)** Graphs showing normalized movement, captured by video synchronized to EEG, as a function of SWD duration (r = - 0.9513; left) or inter-SWD interval (r = 0.9871; right). Pearson correlation test. *All gra*p*hs represent mean ± SEM*. *n.s.: non significant*.

### Excitability phenotypes in *Gnb1*^K78R/+^ mice

We did not visually observe recurrent spontaneous convulsive seizures in *Gnb1*^K78R/+^ mice, nor did we observe sudden death in the colony which might have indicated lethal seizures. We therefore assessed seizure susceptibility by electroconvulsive threshold (ECT) testing in B6NJ mice using stimulus ECT settings for the minimal generalized (clonic forebrain) seizure endpoint. Both female and male *Gnb1*^K78R/+^ mice have a significantly lower ECT compared to WT mice (Figure 2A). Similar results were obtained on the F1H background (Figure 2A).

We further used video-electroencephalogram (EEG) to examine F1H mice for spontaneous seizure events. *Gnb1*^K78R/+^ mice exhibited very frequent generalized bilateral spike-and-wave discharges (SWD) (Figure 2B), between 1,000 and 3,000 SWDs per 24 h, while WT mice exhibited extremely rare SWDs (Figure 2C; mean ± SEM WT: 11.7 ± 6.5 vs. *Gnb1*^K78R/+^: 1723 ± 108.9). The mean duration of SWDs ranged from 2.28s s to 6.74 s (Figure 2D), with some lasting longer than 30 s, and the fundamental frequency that shows peak spectral power was 7.34 Hz (Figure 2E). We observed similarly frequent SWDs on the B6NJ background (data not shown). SWD events were correlated with wake and non-rapid eye movement (NREM) sleep epochs, which revealed a 74% probability to be in wake state (Figure 2F,G; mean ± SEM wake: 0.74 ± 0.01 vs. NREM: 0.075 ± 0.007). Normalized delta power in the pre-SWD period is no different from that in the total wake period, further confirming SWD events predominately occur during wakefulness (Figure 2G). The wake-prevalence of SWDs is not due to the mice spending more time in wake than in NREM sleep. Indeed, over a 24 h period, WT and *Gnb1*^K78R/+^ mice spend a similar amount of time in wake and NREM sleep (Figure 2G), although *Gnb1*^K78R/+^ mice spend slightly more time in NREM sleep than wake, due to increased probability of entering NREM sleep after an SWD event (Figure 2F; blue vs purple lines).

Frequent SWDs in rodent models is indicative of absence seizures (Akman et al., 2010; Kim et al., 2015; Tokuda et al., 2007), which are usually accompanied by behavioral arrest (Frankel et al., 2005; Pearce et al., 2014; Tan et al., 2007). We used a camera synchronized to the EEG with automated tracking to record mouse movement and observed that movement correlated with inter-SWD interval duration, and inversely correlated with SWD duration, demonstrating some behavioral arrest during the longest SWD events (Figure 2H). Together, these data show that *Gnb1*^K78R/+^ mice exhibit very frequent, generalized, non-convulsive absence-like seizures.

### *Gnb1*^K78R/+^ cortical networks display a bursting phenotype

We used multi-electrode arrays (MEA) recordings to characterize the spontaneous activity of *in vitro* neural networks formed from WT and *Gnb1*^K78R/+^ neurons harvested from P0 cortices, and analyzed a variety of spiking and bursting features (see Methods). We show the combination of 9 plates corresponding to 8 biological replicates (Figure 3A). For each feature, we normalized data recorded on each plate by dividing the activity in each well by the mean activity of WT wells. We then performed a Mann-Whitney U (MWU) test followed by 1000 permutations to compare activity in WT versus mutant wells across the time points of interest. Graphs showing the combined analysis without normalization are available in Figure S4E. We also combined the permutated MWU p-values from each individual plate using Fisher’s method, which gave similar results to the WT normalization. The p-values using both methods are available in Table S1.

**Figure 3.**
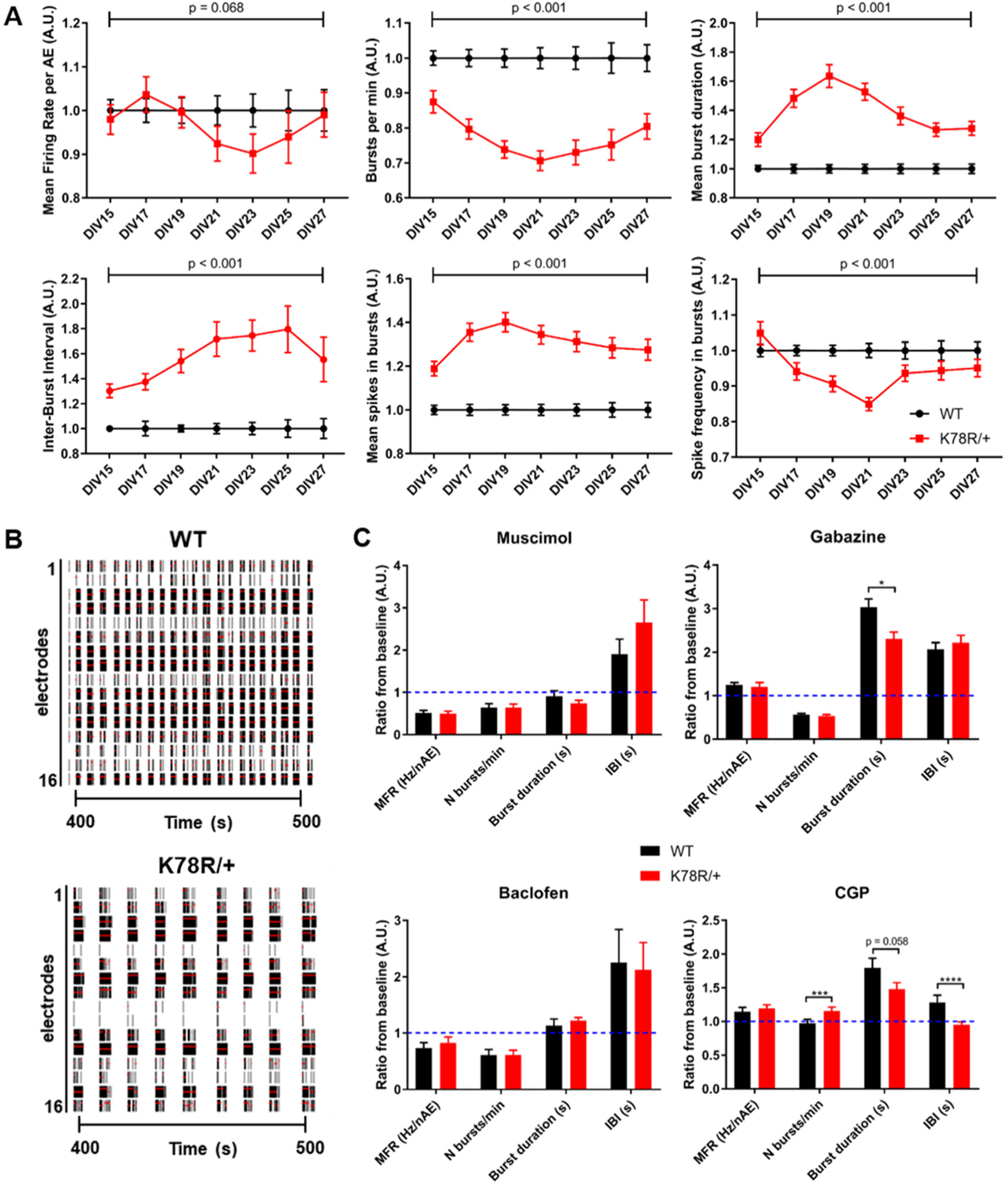
Excitability phenotypes in *Gnb1*^K78R/+^ cortical neurons. **(A)** Graphs representing spontaneous activity on MEA of *Gnb1*^K78R/+^ cortical neurons (red) normalized to WT cortical neurons (black) from DIV15 to DIV27. From left to right: MFR per active electrodes, number of bursts per minute, mean burst duration, mean IBI, mean spikes in bursts and mean frequency of spikes in bursts. n = 144 WT wells and 148 *Gnb1*^K78R/+^ wells from 9 plates (8 biological replicates = 8 independent primary cultures); Permutated p-values calculated with a Mann-Whitney U test followed by 1000 permutations are indicated on the graphs. **(B)** Raster plots showing WT and *Gnb1*^K78R/+^ firing across the 16 electrodes of a well of a representative plate, between 400 and 500 seconds of a 15 min recording at DIV23. The black bars indicate spikes, and the red bars indicate bursts. **(C)** Graphs representing changes in activity of WT and *Gnb1*^K78R/+^ neurons in mature cortical cultures following modulation of GABAA (top panel) and BABAB (bottom panel) receptors. The activity was recorded 10 min after drug application and is normalized to baseline activity before drug application (dashed blue line). MFR, number of bursts per minute, mean burst duration and mean IBI are shown. Muscimol (500 nM; n = 32 WT wells and 29 *Gnb1*^K78R/+^ wells from 4 biological replicates); Gabazine (1 µM; n = 45 WT wells and 40 *Gnb1*^K78R/+^ wells from 5 biological replicates); Baclofen (500 nM; n = 12 WT wells and 11 *Gnb1*^K78R/+^ wells from 1 biological replicate); CGP (1 µM; n = 72 WT wells and 69 *Gnb1*^K78R/+^ wells from 5 biological replicates); Genotype comparisons: *p < 0.05; ***p < 0.001; ****p < 0.0001. Mann-Whitney U test with Bonferroni correction. See also Figure S5. *MFR, mean firing rate; IBI, inter-burst interval*.

By comparing the distribution of mean firing rate (MFR) of WT neurons across DIVs, we observed an increase in MFR until DIV13, followed by stabilization of MFR. We thus present data from DIV15 to DIV27, when the neural networks are established. Both WT and *Gnb1*^K78R/+^ networks developed at the same time, as observed by a similar increase in the number of active electrodes (nAE) per well, reaching a maximum by DIV15 (Figure S4A). We observed a striking bursting phenotype in *Gnb1*^K78R/+^ networks relative to WT. Mutant neurons showed a decrease in burst frequency (decreased number of bursts per minute) as they fired very long bursts followed by very long inter-burst intervals (IBIs), leading to an increase of mean number of spikes in bursts (Figure 3A). We observed only a small and delayed decrease in MFR, suggesting that the MFR is not driving the bursting phenotype; however, there was a decreased spike frequency within bursts (Figure 3A). This bursting phenotype is clearly visible on raster plots (Figure 3B).

At the cell density used here, primary neural networks become highly synchronized over time (Chiappalone et al., 2006; McSweeney et al., 2016; Wagenaar et al., 2006). Therefore, we evaluated key synchrony parameters including: percentage of spikes in network spikes (NS) (Figure S4B), which identifies highly synchronized short events (10 ms), as well as mutual information (Figure S4C) and STTC (Figure S4D), which assess pairwise electrode correlations between adjacent electrodes and at the well-level, respectively (Cutts and Eglen, 2014; Gelfman et al., 2018). Interestingly, we did not observe significant differences in synchrony between WT and *Gnb1*^K78R/+^ networks (Figure S4B-D); however, when we compared the percentage of spikes in NS of our WT cultures with lower density cultures, we still did not observe a major difference (data not shown), suggesting that altered synchrony is not underlying the bursting phenotype *Gnb1*^K78R/+^ networks.

### Differential response of *Gnb1*^K78R/+^ neural networks to GABA_A_ and GABA_B_ receptor modulation

To assess mature network response to inhibition, we acutely treated WT and *Gnb1*^K78R/+^ cortical neurons with muscimol, an agonist of ionotropic GABA_A_ receptors (GABA_A_R) (Cherubini, 2012), or baclofen, an agonist of metabotropic GABA_B_ receptors (GABA_B_R) (Gassmann and Bettler, 2012). Both drugs decreased MFR of WT and *Gnb1*^K78R/+^ neurons, showing that mutant neural networks respond normally to inhibition (Figure 3C). Muscimol had a very potent effect on overall network activity, while baclofen decreased burst frequency and mildly increased burst duration (Figure 3C; Figure S4F,H).

We further treated the cultured networks with the antagonists gabazine for GABA_A_R, or CGP 55845 for GABA_B_R, which are expected to increase excitability. Indeed, gabazine application led to WT network bursting patterns reminiscent of *Gnb1*^K78R/+^ bursting, while it exacerbated the bursting phenotype of *Gnb1*^K78R/+^ networks (Figure 3C; Figure S4G). Together, these data suggest that decreased inhibitory tone in *Gnb1*^K78R/+^ neural networks underlies their aberrant bursting.

While CGP-mediated inhibition of GABA_B_R in WT networks also yielded increased excitability and bursting, *Gnb1*^K78R/+^ networks responded in a significantly differently manner relative to WT. CGP increased burst frequency, while more modestly increasing mean duration than in WT networks, thereby not aggravating *Gnb1*^K78R/+^ aberrant bursting (Figure 3C; Figure S4I). These data suggest that GABA_B_R signaling is affected by the K78R mutation, possibly contributing to the decreased inhibitory tone.

### Pharmacological response of the pre-clinical models

Ethosuximide (ETX) and valproic acid (VPA) are two antiepileptic drugs (AED) commonly used to treat absence seizures in humans, which also interrupt SWDs rodent models (Kim et al., 2015; Manning et al., 2003; Marescaux et al., 1992; Vrielynck, 2013). Acute treatment of *Gnb1*^K78R/+^ mice with either ETX or VPA nearly abolished SWDs (Figure 4A). Owing to the short half-life of both drugs, SWDs progressively recovered an hour after treatment. On the other hand, phenytoin (PHT), an AED with a probable worsening effect on absence seizures in humans and rodents (Kim et al., 2015; Manning et al., 2003; Marescaux et al., 1992; Vrielynck, 2013), did not affect SWD occurrence (Figure 4A).

**Figure 4.**
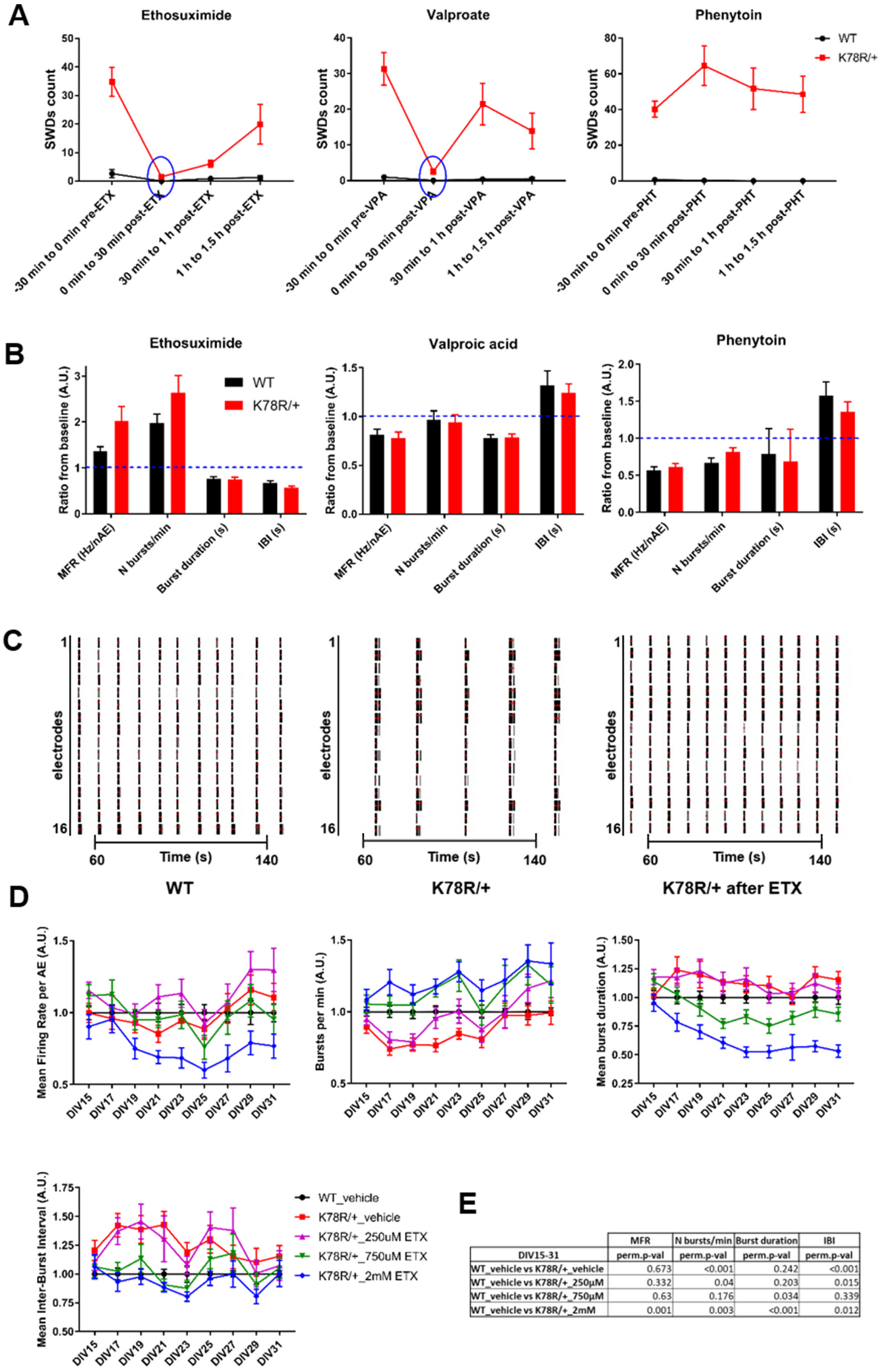
Pharmacological response of the pre-clinical models. **(A)** SWD manual counts on EEG recordings before and after acute treatment of adult mice with the AEDs ethosuximide (ETX), valproate (VPA), and phenytoin (PHT). n = 17 WT and 21 *Gnb1*^K78R/+^ for ETX, 12 WT and 21 *Gnb1*^K78R/+^ for VPA, and 5 WT and 7 *Gnb1*^K78R/+^ for PHT. **(B)** Graphs representing changes in activity of WT and *Gnb1*^K78R/+^ neurons in mature cortical cultures on MEA following modulation by the AEDs ETX, VPA and PHT. The activity was recorded 10 min after drug application and is normalized to baseline activity before drug application (dashed blue line). MFR, number of bursts per minute, mean burst duration and mean IBI are shown. ETX (4 mM; n = 52 WT wells and 51 *Gnb1*^K78R/+^ wells from 5 biological replicates); VPA (1 mM; n = 47 WT wells and 50 *Gnb1*^K78R/+^ wells from 5 biological replicates); PHT (20 µM; n = 28 WT wells and 26 *Gnb1*^K78R/+^ wells from 4 biological replicates). Genotype comparison: n.s. Mann-Whitney U test with Bonferroni correction. **(C)** Raster plots showing WT and *Gnb1*^K78R/+^ firing before and 10 min after acute application of 4mM ETX across the 16 electrodes of a well of a representative plate, between 50 and 150 seconds of a 3 min recording at DIV22. The black bars indicate spikes, and the red bars indicate bursts. **(D)** Graphs representing spontaneous activity on MEA from DIV15 to DIV31 of untreated *Gnb1*^K78R/+^ cortical neurons (red), and *Gnb1*^K78R/+^ cortical neurons treated with 250 µM (purple), 750 µM (green) or 2 mM (blue) ETX chronically from DIV3 to DIV31, normalized to WT cortical neurons (black). MFR, number of bursts per minute, mean burst duration and mean IBI are shown from left to right. n = 30 wells from 5 biological replicates for all genotypes/conditions. **(E)** Table showing the statistics for the graphs shown in D. Mann-Whitney U test with 1000 permutations. *All graphs represent mean ± SEM. DIV: days in vitro. n.s., non significant.* See also Figure S5.

We investigated the effects of these AEDs on neural networks on MEA. Interestingly, ETX, but not VPA, had a rescue effect on the aberrant bursting of *Gnb1*^K78R/+^ networks. ETX increased the MFR by 2-fold and the number of bursts by 2.5-fold, with an apparent stronger effect on *Gnb1*^K78R/+^ neurons than WT neurons, although the difference was not significant (Figure 4B). ETX also decreased the mean duration of bursts and the IBIs (Figure 4B). Overall, ETX restored normal firing to the mutant networks, as can easily be observed on the raster plots before and after drug application (Figure 4C). On the other hand, VPA had a mild, non-specific effect on both WT and mutant networks, dampening overall neuronal activity (Figure 4B) and PHT similarly reduced overall firing (Figure 4B). These observations suggest that ETX mechanism of action is specific for Gβ1 signaling.

To determine whether chronic ETX dosing could have a lasting effect on the phenotype, we applied ETX on the cultured networks at three different doses (250 μM, 750 μM and 2 mM), every other day from DIV3 until DIV31 and observed that ETX reverted *Gnb1*^K78R/+^ aberrant bursting in a dose-dependent manner (Figure 4D,E). At 250 μM, ETX had no effect on the bursting phenotype, while at 750 μM, ETX did restore close to normal firing to the mutant network in a prolonged manner. It increased number of bursts per minute, decreased mean duration of bursts and decreased IBIs, compared to the vehicle- or 250 μM-treated *Gnb1*^K78R/+^ networks. At 2 mM, ETX effect was too strong, greatly reducing network activity. Overall, these data show that while having a short half-life, chronic ETX dosing leads to a cumulative and prolonged rescue of the bursting phenotype.

### GIRK channel dysregulation partially contributes to the bursting phenotype on MEA

Evidence suggests ETX primarily acts through inhibition of low voltage-activated T-Type Ca^2+^ channels (Ca_v_3) (Coulter et al., 1989, 1990), although it has also been shown to inhibit GIRK channels *in vitro* (Kobayashi et al., 2009). We therefore evaluated whether ETX effects on *Gnb1*^K78R/+^ network bursting could be recapitulated by Ca_v_3 or GIRK antagonists.

Two Ca_v_3 antagonists, ML218 (Xiang et al., 2011) and NNC 55-0396 (Huang et al., 2004), were tested on MEA. ML218 decreased mean burst duration and increased mean IBIs of both WT and *Gnb1*^K78R/+^ networks, reducing overall network activity (Figure 5A). NNC 55-0396 showed a different response than ML218, suggesting that the two inhibitors may target different channel subtypes, and had no major effect on the features assessed (Figure 5A). These results suggest that increased Ca_v_3 activity is not a major contributor to the bursting phenotype, and that ETX main mechanism in the cortex may not be inhibition of these channels.

**Figure 5.**
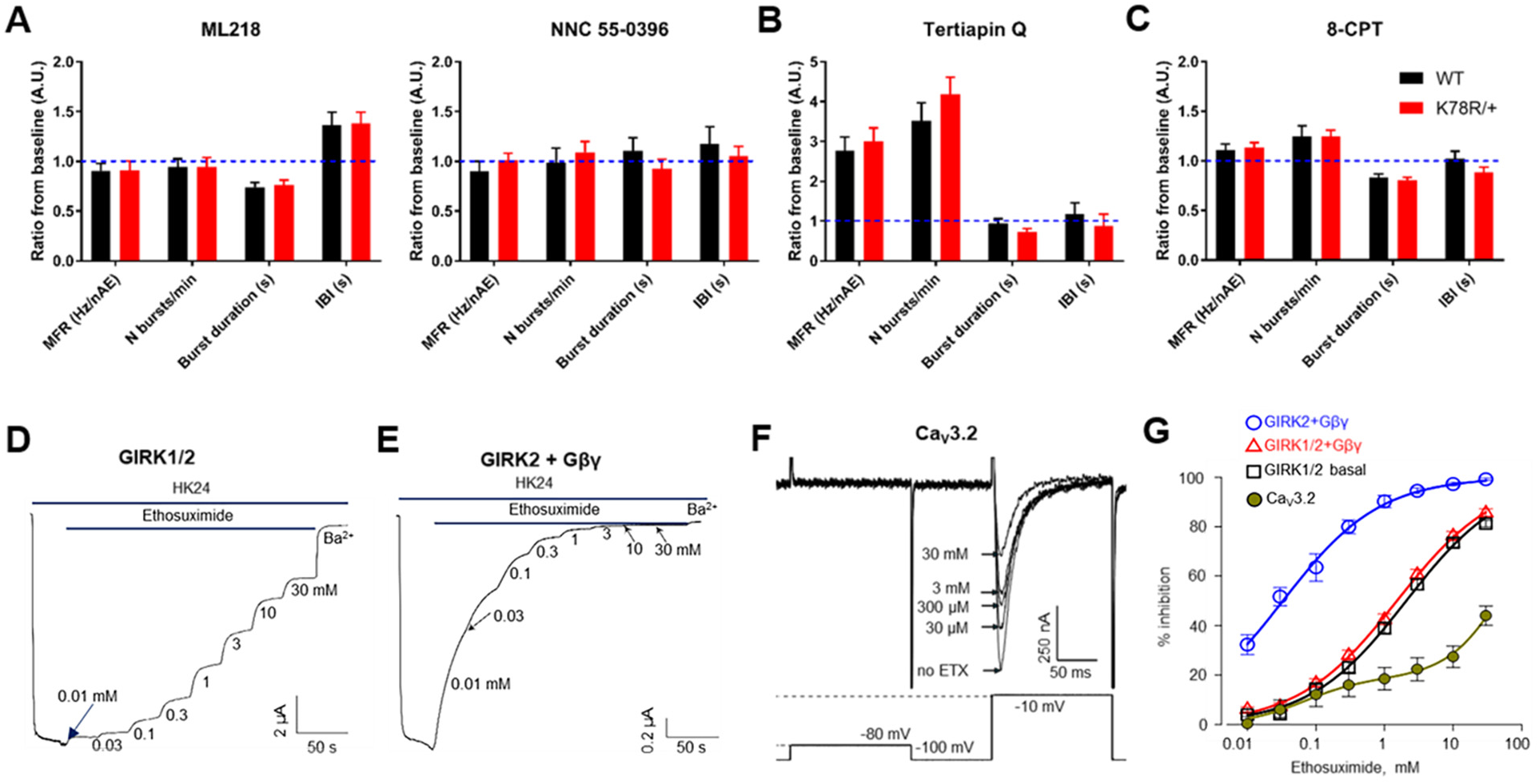
Ethosuximide rescues the bursting phenotype partially through GIRK inhibition. **(A-C)** Graphs representing changes in activity of WT and *Gnb1*^K78R/+^ neurons in mature cortical cultures following modulation of T-type Ca^2+^ channels **(A)**, GIRK channels **(B)**, or A1A receptors **(C)**. The activity was recorded 10 min after drug application and is normalized to baseline activity before drug application (dashed blue line). MFR, number of bursts per minute, mean burst duration and mean IBI are shown. ML218 (750 nM; n = 38 WT wells and 39 *Gnb1*^K78R/+^ wells from 5 biological replicates); NNC 55-0396 (500 nM; n = 31 WT wells and 36 *Gnb1*^K78R/+^ wells from 3 biological replicates Tertiapin Q (10 µM; n = 55 WT wells and 52 *Gnb1*^K78R/+^ wells from 3 biological replicates); 8-CPT (200 nM; n = 40 WT wells and 37 *Gnb1*^K78R/+^ wells from 3 biological replicates). Genotype comparisons: *p < 0.05; Mann-Whitney U test with Bonferroni correction. **(D,E)** Representative record showing the effect of ETX on Ibasal of GIRK1/2 **(D)** and GIRK2 activated by coexpressed Gβγ **(E)**. Sequential application of increasing doses of the drug inhibited the current. 2.5 mM Ba^2+^ was added at the end of protocol for a complete inhibition of GIRK. **(F)** The lower panel shows the voltage protocol used to elicit Ba^2+^ current through T-type CaV3.2 channel, I_Ba_, in the study of dose-dependent effect of ETX. The voltage step from −80 to −100 mV was used to measure leak currents used for the calculation of net IBa. Cav3.2 yielded typical IBa with fast activation and inactivation. The upper panel shows examples of inhibition of IBa by ETX in one cell (4 out of 8 concentrations used are shown). **(G)** Dose-dependence curves of ETX inhibition. Each circle is mean ± SEM, n = 6 to 18. The solid lines show fits to Hill equation or two-site binding equation. *All graphs represent mean ±* SEM. *ETX: ethosuximide.* See also Figure S6.

We next tested Tertiapin Q, a GIRK channel antagonist (Jin et al., 1999; Jin and Lu, 1999). Tertiapin Q ameliorated greatly the *Gnb1*^K78R/+^ bursting phenotype, by strongly increasing MFR (3-fold) and number of bursts per minute (4-fold), while decreasing mean duration of bursts and more modestly mean IBI (Figure 5B; Figure S5A). We previously observed a differential response to GABA_B_R inhibition between *Gnb1*^K78R/+^ and WT neurons (Figure 3C). While GABA activation of GABA_B_R is a major regulator of GIRK channels at inhibitory synapses, the ambient neurotransmitter adenosine, which is thought to participate to tonic inhibition, activates adenosine A_1_ receptors (A_1_AR) which also regulate GIRK channels (Luscher and Slesinger, 2010). We applied a potent A_1_AR antagonist, 8-Cyclopentyl-1,3-dimethylxanthine (8-CPT) (Bruns et al., 1987), and observed a mild rescue effect of the bursting phenotype, through an increase in MFR and number of bursts per minute, and reduction of mean burst duration and mean IBI (Figure 5C; Figure S5B). Altogether, these data point toward Gβγ regulation of GIRK channels as a contributing mechanism to the bursting phenotype of *Gnb1*^K78R/+^ cortical networks, and suggest that ETX may act partially through GIRK inhibition to restore normal firing.

### Ethosuximide differentially inhibits GIRK1/2, GIRK2 and Ca_v_3.2 channels

We further tested the hypothesis that ETX ameliorates the MEA bursting phenotype through GIRK inhibition by assessing ETX inhibition of heterotetrameric GIRK1/2 (overall most abundant GIRK in the mammalian brain (Luscher and Slesinger, 2010)) and homotetrameric GIRK2 (abundant in some of the midbrain nuclei (Lujan and Aguado, 2015)), in *Xenopus* oocytes. Both basal and Gβγ-evoked GIRK1/2 currents were inhibited in a dose-dependent manner by ETX (Figure 5D,G), with an IC_50_ of ∼1.5-2 mM (Figure 5G), similar to a previous report (Kobayashi et al., 2009). Remarkably, Gβγ-activated GIRK2 was inhibited at much lower doses (Figure 5E), with an IC_50_ of ∼30 µM (Figure 5G). I_basal_ of GIRK2 was too small to reliably monitor inhibition, explaining the IC_50_ discrepancy with a previous report (Kobayashi et al., 2009). For comparison, we tested the effect of ETX on Ca_V_3.2, a T-type Ca^2+^ channel abundant in the brain. Expression of Ca_V_3.2 yielded typical transient Ba^2+^ currents (Figure 5F; Figure S5C), which were inhibited by ETX in a dose-dependent manner, but less efficiently compared to GIRK channels, with an apparent IC_50_ of > 30 mM (Figure 5G).

The dose-response curves for ETX inhibition of both GIRK1/2 and GIRK2 were well fitted by a one-site Hill equation with apparent K_d_ (K_d,app_) values close to the observed IC_50_ (Figure 5G; Figure S5D-F). However, the low Hill coefficient (n_H_ ∼0.6) could indicate two binding sites with different affinity - satisfactory fits were also obtained assuming 2 affinity components (Figure S5D) - or two populations of channels with different sensitivity to block. In comparison, for Ca_v_3.2, a very good fit was obtained only with a two-component binding isotherm: a putative high-affinity site with K_d,app_ of 67 µM, however accounting for only < 20% of current inhibition, and a low-affinity site with an estimated K_d,app_ of 69mM (Figure 5G; Figure S5D-F). In summary, both visual examination and results of fitting indicate that, in this model system, GIRK channels, especially GIRK2, are blocked by ETX more effciently than T-type Ca^2+^ channels.

### K78R modulates GIRK1/2 and GIRK2 channel activation

To study the effects of the K78R mutation on GIRK channel activation by Gβγ, GIRK1/2 or GIRK2 channels were expressed in *Xenopus* oocytes together with Gβ_WT_γ or Gβ_K78R_γ (Rubinstein et al., 2009). Exchange from physiological low-K^+^ ND96 to high-K^+^ HK24 solution resulted in the development of inward currents carried mostly by GIRKs (Figure 6A,F). GIRK1/2 and GIRK2 show substantial differences in gating: GIRK1/2 shows high, Gβγ-dependent basal current (I_basal_) and a modest 1.5-4 fold activation by Gβγ (Figure 6A; compare black and dark green traces), whereas GIRK2 has low, Gβγ-independent I_basal_ and shows strong ∼30-50 fold activation by Gβγ (Dascal and Kahanovitch, 2015) (Figure 6F; compare black and dark green traces).

**Figure 6.**
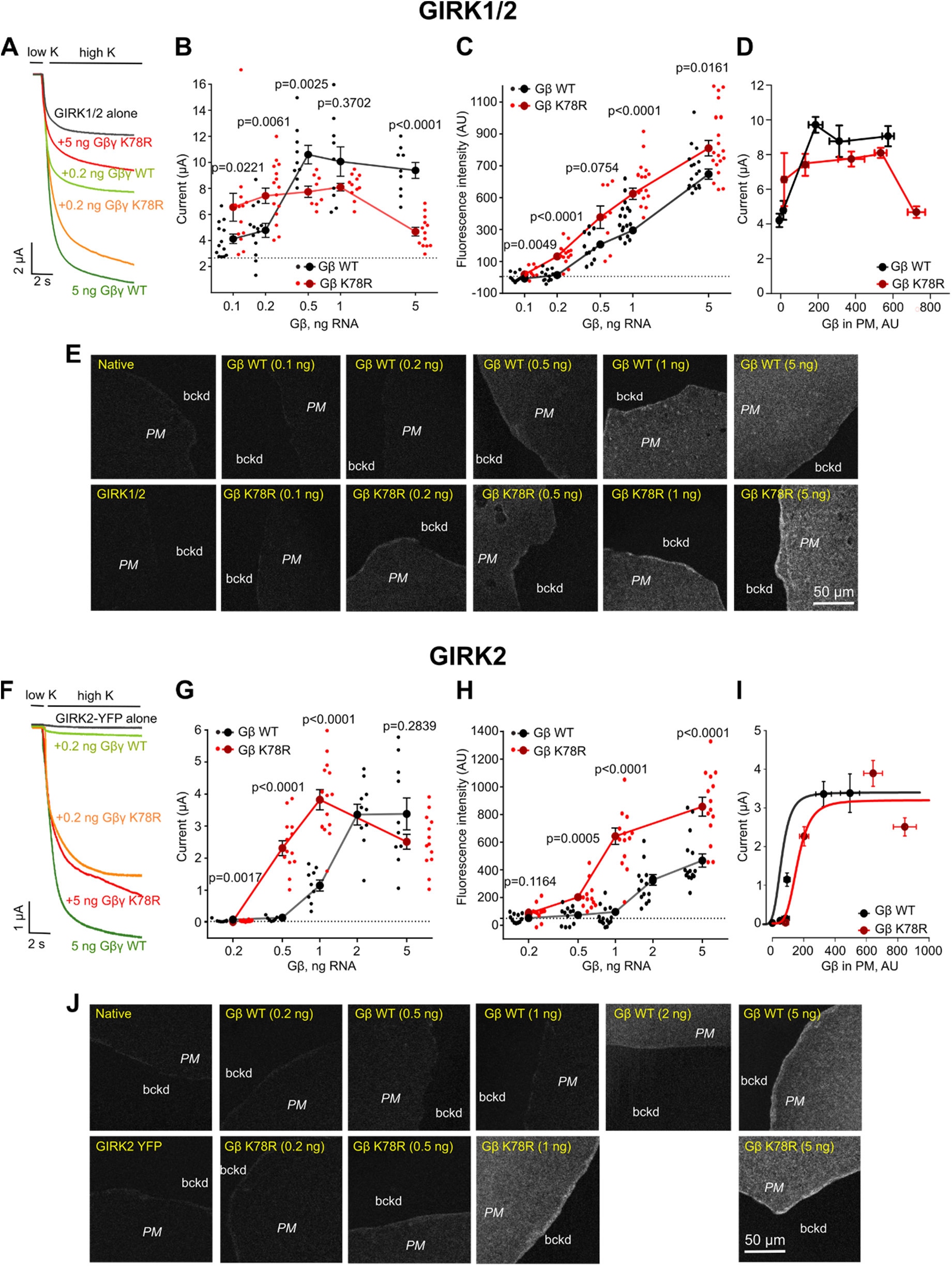
K78R is a GoF for GIRK1/2 and GIRK2 activation but, at high expression levels, shows a LoF for GIRK1/2. **(A)** Representative records of GIRK1/2 currents in *Xenopus* oocytes injected with RNAs of GIRK1 and GIRK2 (0.05 ng each), and the indicated amounts of Gβ RNA. The amount of Gγ RNA was 1/5 of Gβ. **(B)** Summary of GIRK1/2 activation by Gβ_WT_γ and Gβ_K78R_γ in the experiment shown in A. Small circles represent one cell and large circles show mean ± SEM of total GIRK1/2 current for each Gβ RNA dose. The dotted line shows the basal level of current in oocytes injected with GIRK1/2 RNA without coexpression of Gβγ. For each Gβ dose, raw data for Gβ_WT_ and Gβ_K78R_ were compared using two-tailed t-test or Mann-Whitney rank sum test (see Methods for details). **(C)** Changes in surface expression of Gβ_WT_ and Gβ_K78R_ as a function of RNA dose, in the presence of GIRK1/2, measured in GMP. Data shown are the net fluorescent signal produced by the expressed Gβ, after subtraction of the average signal observed in naïve (uninjected) oocytes. **(D)** Dose-dependent activation of GIRK1/2 vs. actual surface expression of Gβγ as measured in GMP. **(E)** Representative images of membrane patches at different Gβ RNA doses in the presence of GIRK1/2. **(F)** Representative records of GIRK2 currents in *Xenopus* oocytes injected with RNAs of GIRK2 (2 ng; C-terminally YFP-labeled GIRK2 was used in this experiment) and the indicated amounts of Gβ RNA. The amount of Gγ RNA was 1/5 of Gβ. **(G)** Summary of the experiment shown in F. **(H)** Changes in surface expression of Gβ_WT_ and Gβ_K78R_ as a function of RNA dose, in the presence of GIRK2, measured in GMP. Analysis was done as in C. **(I)** Dose-dependent activation of GIRK2 vs. actual surface expression of Gβγ as measured in GMP. The results were fitted to Hill equation with a fixed Hill coefficient of 4. The fit yielded dissociation constant (Kd) of 118 and 162 AU and maximal currents of 3.4 and 3.18 µA for Gβ_WT_γ and Gβ_K78R_γ, respectively. **(J)** Representative images of membrane patches at different Gβ RNA doses in the presence of GIRK2. See also Figure S6. *GMP, giant excised plasma membrane patches; AU, arbitrary units*.

To capture the dose dependency of Gβγ activation on GIRK channels, we titrated the amount of expressed Gβγ protein by injecting increasing amounts of Gβ_WT_ or Gβ_K78R_ and Gγ RNA (5/1 RNA amounts, w/w). For GIRK1/2, small activation by Gβ_WT_γ was observed with 0.1 ng Gβ RNA, and saturation was reached at 0.5-1 ng Gβ RNA (Figure 6B and Figure S5G). Interestingly, at low doses of Gβ RNA (0.05-0.2 ng), Gβ_K78R_γ activated GIRK1/2 better than Gβ_WT_γ (Figure 6A,B and Figure S5G), suggesting gain-of-function (GoF). However, at higher Gβ doses, Gβ_K78R_γ ability to activate GIRK1/2 progressively decreased, until being almost completely lost with 5 ng Gβ RNA (Figure 6B and Figure S5G).

To investigate whether this loss-of-function (LoF) was due to poor surface expression of Gβ_K78R_, we monitored the levels of membrane-attached Gβγ in giant excised membrane patches (GMP) of oocytes (Figure 6E) (Peleg et al., 2002; Rubinstein et al., 2009). We found that Gβ_K78R_γ consistently showed higher surface density than Gβ_WT_γ at all RNA doses (Figure 6C). To directly compare the actual dose-dependence of GIRK1/2 activation for Gβ_WT_γ and Gβ_K78R_γ, we plotted GIRK1/2 currents as a function of Gβγ surface levels as measured in oocytes of the same groups (Figure 6D). Per equal amount of expressed Gβγ, Gβ_K78R_γ activated GIRK1/2 similarly, or even less, compared to Gβ_WT_γ. The highest dose of Gβ_K78R_γ caused a large decrease in GIRK1/2 current, but we did not reach a similar expression level of Gβ_WT_γ to allow a direct comparison. The tendency for overall smaller maximal activation of GIRK1/2 by Gβ_K78R_γ was consistently observed in additional experiments (Figure S6F). In all, these results suggest that K78R is a gain-of-expression mutation, and the apparent GoF is due exclusively to higher surface expression compared to Gβ_WT_γ. In fact, the smaller maximal activation and the strong attenuation of current by high doses of Gβ_K78R_γ indicated that K78R may be a partial LoF for GIRK1/2.

The activation of GIRK2 by coexpressed Gβγ showed a similar pattern, with an apparent strong GoF for Gβ_K78R_γ at lower Gβ RNA doses which subsided at saturating RNA doses, 2-5 ng (Figure 6F,G and Figure S5H). With 0.5 ng Gβ RNA, GIRK2 current evoked by Gβ_K78R_γ was more than 6-fold larger than with Gβ_WT_γ (Figure 6G). Like with GIRK1/2, the surface expression of Gβ_K78R_γ was higher than Gβ_WT_γ at all RNA doses (Figure 6H,J). Re-plotting GIRK2 activation as a function of Gβγ surface levels showed that K78R activated GIRK2 similarly to Gβ_WT_γ in the range of surface expression studied (Figure 6I). Thus, similarly to GIRK1/2, the physiological GoF of Gβ_K78R_γ is due to gain of expression. In conclusion, altered GIRK channel activation by the K78R mutation is likely contributing to disease.

## DISCUSSION

We describe the first mouse model of GNB1 Encephalopathy, carrying the pathogenic missense variant K78R. Prior studies emphasized the essential role of Gβ1 during brain development, as evidenced by the embryonic lethality of *Gnb1* knockout mice due to neural tube closure and neuronal progenitor cell proliferation defects (Okae and Iwakura, 2010). Here, we show that while the heterozygous K78R mutation does not impair brain development to the same degree as the knockout, it does affect neuronal activity by altering Gβ1 modulation of ion channels essential for the regulation of neuronal excitability.

*Gnb1*^K78R/+^ mice phenocopy the main aspects of GNB1 Encephalopathy (Hemati et al., 2018), including developmental delay, and motor and cognitive deficits. Nearly 60% of GNB1 patients have an abnormal EEG, suggesting that *GNB1* mutations confer abnormal neuronal excitability. Our studies revealed that *Gnb1*^K78R/+^ mice are not only more susceptible to generalized seizures, but also have a very high incidence absence-like seizures. Evaluation of mature neuronal network properties using MEA further revealed that *Gnb1*^K78R/+^ cortical neurons have a hyperexcitability phenotype, firing long bursts interspaced with long quiet intervals. Activity-dependent network plasticity during early development is essential to establish precise connections for proper adult circuit activity, a process in which GABA signaling plays a major role (Ben-Ari, 2001; Ben-Ari et al., 2012a; Cellot and Cherubini, 2013; Kirkby et al., 2013). Indeed, GABA exerts an early developmental depolarizing effect that progressively becomes hyperpolarizing, towards the beginning of the second postnatal week in rodents, and around DIV11 *in vitro* (Ben-Ari et al., 2007; Ben-Ari et al., 2012b; Ganguly et al., 2001; Leonzino et al., 2016; Tyzio et al., 2007; Valeeva et al., 2016). To this point, modulation of GABA_A_ signaling on MEA suggests a reduced inhibitory tone in *Gnb1*^K78R/+^ networks. An impaired GABA switch is thought to underlie several neurodevelopmental disorders (Amin et al., 2017; He et al., 2014; Lysenko et al., 2018), as well as abnormal hippocampal activity in the *tottering* mouse model of absence epilepsy (Nakao et al., 2015), and thus could possibly underlie *Gnb1*^K78R/+^ network deficits.

It is well established that SWDs result from hypersynchronized oscillations in the cortico-thalamo-cortical networks. These oscillations were commonly thought to originate from sustained burst firing of thalamic neurons (Danober et al., 1998; Huguenard and McCormick, 2007), underlain by increased Ca^2+^ currents through T-type voltage-gated calcium channels (Cain and Snutch, 2013; Cain et al., 2018; Cheong and Shin, 2013). However, there is compelling evidence for a focal origin in specific regions of the somatosensory cortex (McCafferty et al., 2018; Meeren et al., 2002; Polack et al., 2007; Zheng et al., 2012). Interestingly, ETX is able to suppress SWDs originating in the cortical focus and restore normal neuronal activity (Manning et al., 2004; Polack and Charpier, 2009). We determined that VPA and ETX transiently abolished the SWDs in *Gnb1*^K78R/+^ mice; however, only ETX restored normal bursting of cortical neural networks *in vitro*. Because the cultured neuronal networks lack some of the disease mechanism-independent circuits through which AEDs act and yet they reveal clear effects of the mutation, this suggests the possibility of mechanistic specificity of ETX towards the Gβ1 signaling affected by K78R. ETX is thought to act mainly as a T-type Ca^2+^ channel blocker, although controversy remains as to whether this block occurs at physiologically relevant doses (Crunelli and Leresche, 2002; Gomora et al., 2001; Todorovic and Lingle, 1998). ETX may reduce the Ca^2+^-activated K^+^ current and the persistent Na^+^ current (Broicher et al., 2007; Leresche et al., 1998), and was shown to inhibit basal GIRK currents in *Xenopus* oocytes (Kobayashi et al., 2009).

To gain further insight, we compared the inhibitory effects of ETX on GIRK and T-Type Ca^2+^ channels by heterologous expression of GIRK1/2, GIRK2 or Ca_v_3.2 channels in *Xenopus* oocytes. In concordance with a previous report (Gomora et al., 2001), we observed that the effect of ETX on Ca_v_3.2 is state-dependent with the presence of a high-affinity block of ∼20% of inactivated channels, and a low-affinity block for the non-inactivated channels, which are the majority at the holding potentials used (−80 or −100 mV). Overall, ETX appeared to be more efficient in blocking GIRK channels than T-type channels, and blocked GIRK2 homomers much more potently than GIRK1/2 heteromers. Nevertheless, we cannot exclude that at physiological resting potentials of −60 to −70 mV, a larger proportion of T-type channels would be inactivated, increasing the sensitivity to ETX. However, there is no evidence yet supporting a regulation of T-type channels by Gβ1 signaling - Gβ2, but not Gβ1, interacts with Ca_v_3.2 channels (Wolfe et al., 2003) - and inhibition of Ca_v_3 channels on MEA did not improve the bursting phenotype, arguing against the ETX rescue of mutant phenotypes *in vitro* and *in vivo* being due to inhibition of these channels. We observed that inhibition of GABA_B_R or A_1_AR signaling, which both modulate GIRK activation directly through Gβγ, and direct inhibition of GIRK channels, all improved the bursting phenotype. These results collectively implicate increased GIRK signaling as an underlying disease mechanism in GNB1 Encephalopathy.

GIRK channels are essential regulators of neuronal excitability both presynaptically, by inhibiting neurotransmitter release, and postsynaptically by generating slow iPSPs (Nicoll, 1988; Thompson et al., 1993; Wu and Saggau, 1997). While it may seem counterintuitive that blocking GIRK channels would have an antiepileptic effect, there is evidence for the involvement of GABA_B_ signaling in absence seizures. In rodent models, activation of GABA_B_R exacerbates SWDs while inhibition, in either the thalamus or the cortex, suppresses SWDs (Bortolato et al., 2010; Hosford et al., 1992; Manning et al., 2004; Richards et al., 2003). Indeed, it is thought that the GABA_B_R-mediated K^+^ slow IPSPs (generated by GIRK channels) activate the low threshold T-type Ca^2+^ current in thalamocortical cells, priming them for burst firing (Crunelli and Leresche, 1991), supporting the idea that increased GIRK signaling could underlie the bursting phenotype. We could also imagine an antiepileptic effect of GIRK inhibition if the target channels are mainly expressed in interneurons, although it is still unclear which GIRK subunit composition plays a major functional role in these neurons.

We show in *Xenopus* oocytes that Gβ_K78R_ exerts a GoF towards GIRK1/2 and GIRK2 activation at mild Gβ RNA doses, owing to higher surface expression in GMP of Gβ_K78R_ compared with Gβ_WT_, at all RNA doses tested (Dascal, 1997; Peleg et al., 2002; Singer-Lahat et al., 2000). However, at least in the case of GIRK1/2, there appears to also be a change in gating as indicated by the complex dose dependence of GIRK1/2 regulation by Gβ_K78R_γ. Whereas Gβ_K78R_γ showed GoF toward GIRK2 at all Gβγ doses, there was a significant LoF toward GIRK1/2 at high doses, despite robust surface expression of Gβ_K78R_. Further study will be needed to understand this complex regulation. One possibility is a decrease in GIRK function caused by a desensitization-like process caused by Gβγ itself (Ivanova-Nikolova et al., 1998). Indeed, we often observed a maximal Gβ_WT_γ-evoked activation of GIRK1/2 at intermediate RNA doses, followed by a mild decrease at higher doses. This phenomenon could be exacerbated with Gβ_K78R_γ owing to its higher expression. However, the overall lower maximal GIRK1/2 currents observed with Gβ_K78R_γ compared with Gβ_WT_γ at optimal expression levels indicate that Gβ_K78R_ could be a partial LoF mutant toward GIRK1/2. The stronger activation of GIRK1/2 seen with low doses of expressed Gβγ could be a combination of the higher expression of Gβ_K78R_ and the complex, cooperative character of GIRK gating by Gβγ (Ivanova-Nikolova et al., 1998; Sadja et al., 2002; Wang et al., 2016; Yakubovich et al., 2015).

The Gβγ binding sites are located at the interface between two adjacent GIRK subunits (Mahajan et al., 2013; Whorton and MacKinnon, 2013). The discordance of the effect of the K78R mutation on Gβγ activation of GIRK1/2 and GIRK2 suggests previously unsuspected differences in Gβγ-binding surfaces on channels of variable subunit combinations. Further, it suggests that the severity of functional impairment caused by this *GNB1* mutation may differ in neurons with prevalence of GIRK1/2 heterotetramers (e.g. hippocampus) vs. GIRK2 homotetramers (e.g. midbrain) (Lujan and Aguado, 2015; Luscher and Slesinger, 2010).

We assume that mild levels of Gβγ mimic physiological conditions better than the high levels usually attained by saturating expression of a protein in heterologous cell models (Falkenburger et al., 2010; Yakubovich et al., 2015). Therefore, we propose that, under most physiological conditions, K78R is a GoF mutation for GIRK1/2 and GIRK2. Consequently, given the strong inhibitory activity of ETX on GIRK channels, in particular GIRK2 homomers, we hypothesize that ETX exerts its positive effects on cortical neuron bursting, and SWD control, partially through inhibition of the GIRK channel GoF induced by the K78R mutation in the cortex.

In conclusion, our results indicate that the K78R mutation identified in individuals with GNB1 Encephalopathy affects cortical network activity, and suggest that an underlying mechanism is a GoF of GIRK activation. We provide evidence for a specific mode of action of the AED ethosuximide towards the Gβ1 signaling affected by the mutation in the cortex, i.e. inhibition of GIRK channels. Overall, we present the first model of GNB1 Encephalopathy, recapitulating many clinical features and thereby providing an effective preclinical model, and we implicate GIRK channels as a key component of how GNB1 mutations cause disease.

## ACKNOWLEDGMENTS

We thank Dr. Chyuan-Sheng (Victor) Lin of the Transgenics Mouse Shared Resources at Columbia University Irving Medical Center for his help generating the *Gnb1*^K78R/+^ mice. We thank the members of the Institute for Genomic Medicine’s functional groups for helpful discussions, comments and for support, in particular Sarah Dugger. We thank Dr. Tal Keren Raifman for instruction and participation in preparation of giant membrane patches of oocyte membranes. This work was supported by: Israel-India Binational grant ISF_2255_2015 (N.D. and A.K.B); Israel Science Foundation ISF_1282_2018 (N.D.); Grant NIH R37 NS031418 (W.N.F.).

## AUTHOR CONTRIBUTIONS

Conceptualization: S.C., M.J.B., N.D. and D.B.G.; Methodology: S.C., B.S., S.T., M.J.B. and Y.P.; Software: R.S.D., S.G. and Y.P.; Formal Analysis: S.C., B.S., H.P.R., R.S.D., S.G., E.R., Y.P., W.N.F. and D.B.G.; Investigation: S.C., S.P., B.S., H.P.R., G.T., S.T., E.R., Y.P. and W.N.F.; Resources: G.T. and W.N.F.; Writing – Original Draft: S.C.; Writing – Review & Editing: S.C., M.Y. M.J.B., Y.P., W.N.F., N.D. and D.B.G.; Visualization: S.C., B.S., H.P.R. and N.D.; Supervision: A.K.B., M.Y., M.J.B., Y.P., W.N.F., N.D. and D.B.G.; Funding Acquisition: A.K.B., N.D., W.N.F. and D.B.G.

## METHODS

### Animals

#### Mice

*Gnb1*^K78R/+^ were generated using CRISPR/Cas9 at the Transgenics Mouse Shared Resources at Columbia University Irving Medical Center on a C57BL/6NCrl background. Mice were further backcrossed to C57BL/6NJ mice (JAX stock # 005304). For certain experiments, as mentioned in the text, F1 Hybrid mice obtained from a C57BL/6NJ x FVB.129P2 (JAX stock # 004828) mating, were used. Wild-type (WT) littermates were used as controls. Mice were maintained in ventilated cages at controlled temperature (22–23°C), humidity ∼60%, and 12h:12h light:dark cycles (lights on at 7:00AM, off 7:00PM). Mice had access to regular chow and water, *ad libitum.* Mice were bred and procedures were conducted in the Columbia University Institute of Comparative Medicine, which is fully accredited by the Association for Assessment and Accreditation of Laboratory Animal Care, and were approved by the Columbia Institutional Animal Care and Use Committee.

#### Xenopus laevis

Experiments were approved by Tel Aviv University Institutional Animal Care and Use Committee (permits M-08-081 and M-13-002). All experiments were performed in accordance with relevant guidelines and regulations. *Xenopus laevis* female frogs were maintained and operated as described (Dascal and Lotan, 1992; Hedin et al., 1996). Frogs were kept in dechlorinated water tanks at 20±2°C on 10 h light/14 h dark cycle, anesthetized in a 0.17% solution of procaine methanesulphonate (MS222), and portions of ovary were removed through a small incision in the abdomen. The incision was sutured, and the animal was held in a separate tank until it had fully recovered from the anesthesia and then returned to post-operational animals’ tank. The animals did not show any signs of post-operational distress and were allowed to recover for at least 3 months until the next surgery. Following the final collection of oocytes, after 4 surgeries at most, anesthetized frogs were killed by decapitation and double pithing.

### Genotyping

DNA was extracted from tail or ear tissue, and PCR was performed, using the KAPA Mouse Genotyping Standard Kit (KAPA Biosystems). The following primers were used for PCR. Fwd: CGAGCATTGAGATCCTCTTTCT; Rev: GTCATCATTGCTCCATCAACAG. The restriction enzyme *HinfI* was used to distinguish WT from *Gnb1*^K78R/+^ mice.

### DNA constructs and RNA

GIRK1 (rat), GIRK2A (mouse), Gβ1 (bovine) and Gγ2 (bovine) cDNAs used in this study were cloned into high expression oocyte vectors pGEM-HE or pGEM-HJ as described previously 1,4. α1H (human) in pGEM-HEA 11 was kindly provided by Prof. Edward Perez-Reyes. RNA was transcribed in vitro, as described previously 4. The amounts of RNA that were injected, per oocyte: GIRK1/2 ± Gβγ: 0.025-0.5 ng GIRK1 and GIRK2, 5 ng Gβ1, 2 ng Gγ2. GIRK2 ± Gβγ: 2-5 ng GIRK2, 5 ng Gβ1, 2 ng Gγ2. Cav3.2: 5 α1H.

### Protein extraction

P0 cortices were snap-frozen in liquid nitrogen and powdered using a mortar and pestle. Crude protein extracts were prepared by homogenizing the cortices in cold RIPA buffer (Sigma-Aldrich #R0278) containing a protease inhibitor cocktail (Thermo Fisher Scientific #A32955) and a phosphatase inhibitor cocktail (Sigma-Aldrich #4906837001). After vortexing for 1 min, the samples were incubated on a rotator for 15 min at 4°C then centrifuged at 11000 rpm for 15 min at 4°C to collect the supernatant.

For crude membrane/cytosol fractioning, cortices were homogenized in the following homogenization buffer: 0.32 M sucrose, 10 mM HEPES pH 7.4, 2mM EDTA in H2O, containing a protease inhibitor cocktail and a phosphatase inhibitor cocktail. The sampled were homogenized with a motor-driven homogenizer and centrifuged at 1000 g for 15 min at 4°C to remove the pelleted nuclear fraction. The supernatant was further centrifuged at 16000 rpm for 20 min at 4°C to yield the cytosolic fraction in the supernatant and the pelleted membrane fraction. The pellet was resuspended in homogenization buffer. Protein concentrations were determined with the Pierce BCA Protein Assay kit (Thermo Fisher Scientific #23225) using BSA as a standard.

### Western blot

Sample proteins (10 μg) were mixed with 4x LDS buffer (Nupage), and 10x sample reducing agent (Nupage), heated for 10 min at 70°C, separated on 4-12% Bis-Tris mini gel (Nupage) and transfered to a PVDF membrane (Millipore #ISEQ00010). Nonspecific binding was blocked for 1 h at room temperature with 5% blotto (Biorad) in Tris-buffered saline with 0.1% Tween (TBST). Membranes were incubated overnight at 4°C with the primary antibody GNB1 at 1/5000 (Genetex #GTX114442), washed 3x 10 min with TBST and incubated with the secondary antibody HRP-conjugated anti-rabbit at 1/10000 (Invitrogen #32260) in 5% blotto for 1 hour at RT and washed again 3x 10 min with TBST. Blots were incubated for 5 min with Pierce ECL Western Blotting substrate (Thermo Fisher Scientific #32106) and developed with a Kodak X-OMAT 2000A Processor. Blots were stripped for 15 min at RT in ReBlot Plus Strong solution (Millipore #2504), blocked for 30 min in 5% blotto, incubated for 1 hour at RT with an HRP-conjugated β-actin antibody at 1/5000 (Santa Cruz #sc-47778) in 5% blotto and developed as previously described. β-actin was used as a loading control.

### Immunocytochemistry

Primary cortical neurons were seeded on poly-D-lysine-coated 12 mm coverslips in 24-well plates at a density of 150,000 cells/well. At DIV14, cells were quickly washed with phosphate buffered saline (PBS) then fixed with 4% paraformaldehyde for 30 min at RT. Following three washes in PBS, cells were incubated with blocking solution (5% normal donkey serum, 1% bovine serum albumin, 0.3% Triton X-100 in PBS) for 1 hour at RT. Cells were incubated with primary antibodies diluted in blocking solution for 1.5 hours at RT, washed three times with 0.2% Triton X-100 in PBS, then incubated with the secondary antibodies diluted in blocking solution for 30 min at RT in the dark. Finally, the cells were washed two times with 0.2% Triton X-100 in PBS, one time with PBS, and coverslips were mounted on SuperFrost microscopy slides with one drop of Prolong Antifade DAPI (Invitrogen) and allowed to cure overnight at RT in the dark before imaging. Primary antibodies: rabbit anti-GNB1 at 1/200 (Genetex # GTX114442), mouse anti-MAP2 at 1/500 (Sigma # M4403), mouse anti-GFAP at 1/300 (Cell Signaling # 3670), mouse anti-VGlut1 at 1/100 (Synaptic Systems # 135511), mouse anti-Gad67 at 1/500 (EMD # MAB5406), mouse anti-Satb2 at 1/100 (Abcam # ab51502), rat anti-Ctip2 at 1/125 (Abcam # ab18465), rabbit anti-Tbr1 at 1/100 (Abcam # ab31940), mouse anti-Ankyrin G at 1/100 (Santa Cruz # sc-12719). Secondary antibodies: Alexa Fluor 488, 568 or 647 conjugated donkey anti-mouse, donkey anti-rabbit or goat anti-rat were used at 1/500 (Invitrogen). Imaging was performed with an inverted Zeiss AxioObserver Z1 epifluorescent microscope equipped with an Axiocam 503 mono camera, and images acquired with the Zen 2 software. Post-processing was performed with Zen 2 and ImageJ softwares.

### Electroconvulsive threshold (ECT)

Tests were performed in 6-8 weeks old mice using transcorneal electrodes with the Ugo Basile Model 7801 electroconvulsive device as described previously with minor modifications (Frankel et al., 2001). Stimulator settings were 299 Hz, 1.6 ms pulse width, 0.2 s duration, and stimulus was described as iRMS (Integrated root mean square, or area under the curve) calculated from the parameters as: Hz^0.5^ x current x pulse width x duration.

### Electroencephalogram (EEG)

For initial EEG screening, electrode implantation of adult mice approximately 6-8 weeks old and video-EEG was performed essentially as described (Asinof et al., 2016). Mice were anesthetized with tribromoethanol (250 mg/kg i.p., Sigma Aldrich cat# T48402). Three small burr holes were drilled in the skull (1 mm rostral to the bregma on both sides and 2 mm caudal to the bregma on the left) 2 mm lateral to the midline. One hole was drilled over the cerebellum as a reference. Using four teflon-coated silver wires soldered onto the pins of a microconnector (Mouser electronics cat# 575-501101) the wires were placed between the dura and the brain and a dental cap was then applied. The mice were given a post-operative analgesic of carprofen (5 mg/kg subcutaneous Rimadyl injectable) and allowed a 48-hr recovery period before recordings were taken. To record EEG signals, mice were connected to commutators (PlasticOne) with flexible recording cables to allow unrestricted movements within the cage. Signal (200 samples/s) was acquired either on a Grael II EEG amplifier (Compumedics) or on a Natus Quantum amplifier (Natus Neuro), and and the data was analyzed either with the Profusion 5 (Compumedics) or NeuroWorks (Natus Neuro) softwares. Differential amplification recordings were recorded pair-wise between all three electrodes, as well as referential, providing a montage of 6 channels for each mouse. Mouse activity was captured simultaneously by video monitoring using a Sony IPELA EP550 model camera, with an infrared light to allow recordings in the dark. For initial screening and SWD event scoring, we recorded continuously for 24 to 48 hours. For drug dosing studies, mice were recorded continuously for 48 hours, injected with saline around 12pm on day 1 and injected with the drug around 12pm on day 2.

For sleep analysis, we supplemented manual seizure annotation with an automated analysis performed using an in-house developed python script. The script first used fast Fourier transform (FFT) to calculate the power spectrum of the EEG using a 1-sec sliding window, sequentially shifted by 0.25-sec increments. Then it used KMeans clustering algorithm (from scikit-learn library) to generate three clusters based on the distances among delta (1-4 Hz), theta (6-9 Hz) and “seizure”-power (19-23 Hz). We chose the 19-23 Hz band to detect SWD based on its clear separation from normal brain oscillatory activities, although the primary spectral band of SWD in mice is around 7 Hz (overlapped with theta oscillation). The cluster with the highest “seizure”-power was classified as SWD; the cluster with the highest delta power was classified as NREM sleep. The remaining cluster was classified as wake. After clustering, SWD events were further refined by removing false positive events if the averaged “seizure”-power of a given event was lower than two standard deviations from the baseline. Two SWD events were merged into one event if their interval was shorter than 1 sec. Any SWD event with duration less than 0.5 sec was removed for analysis. NREM sleep epochs were refined by removing epochs if their delta power was lower than one standard deviation from the baseline and then by smoothing with a 20-sec sliding window. To correlate SWD events with sleep/wake states, each SWD event was aligned to its onset and the probability of brain states 1 min before and 1 min after event onset was calculated. Notably there are many SWD events in the pre-1 min time window. To minimize the effect of these SWD events on spectral analysis, we calculated the delta power 3 sec prior to the SWD onsets in Figure 2E (pre-SWD). REM could be difficult to distinguish from wake in the absence of EMG signals and was thus combined with wake states. In a few mice with both EEG and EMG recording, we observed about 80 min of REM sleep in Gnb1 mutant mice over a 24-hr period (not shown), which is similar to that in WT mice.

To analyze the effect of SWD events on movement, a custom-built Matlab software was developed to perform real-time video-tracking while simultaneously conducting EEG recording on a Neuralynx Digital Lynx system. The software synced video-taping with EEG recording through a NetCom API provided by Neuralynx. An infrared camera was used to track the body position (the center of the whole body) of a mouse by subtracting each video frame from the background image, captured in the absence of the mouse. The animal’s movement was calculated as the pixel distance between body positions dividing by the time. Then, movement during SWD periods (detected by the algorithm described above) was averaged for each SWD duration or each inter-SWD interval, and further normalized for each animal to the average movement over the whole recording session.

### Multi-electrode arrays (MEA)

#### Plate preparation

One to seven days before dissection, 48-well MEA plates (Axion Biosystems #M768-KAP-48) were coated with 50 µg/mL poly-D-lysine (Sigma-Aldrich #P0899-50MG) in borate buffer, then washed three times with phosphate buffered saline (PBS) and stored in PBS at 4°C until use. Prior to use, PBS was aspirated and plates were allowed to dry in a sterilized hood.

#### Primary neuron culture

Cortical neurons were dissociated from the brains of postnatal day 0 (P0) C57BL/6NJ WT or *Gnb1*^K78R/+^ mice. Pups were immediately decapitated, weighed and genotyped. The entire cerebral cortex was rapidly dissected and cut into small pieces under sterile conditions in cold Hibernate A solution (Gibco, # A1247501). Cortices from two WT or *Gnb1*^K78R/+^ pups were pooled together. The dissected cortices were then enzymatically digested in 20 U mL^-1^ Papain plus DNAse (Worthington Biochemical Corporation, #LK003178 and #LK003172) diluted in Hibernate A for 20 minutes at 37°C. Cells were pelleted by centrifugation at 300RCF for 5 minutes, then the digestion was neutralized by aspirating off the supernatant and adding warm Hibernate A media. Cells were mechanically dissociated by trituration, and counted using a hemocytometer with Trypan blue counterstain. Cells were pelleted by centrifugation at 300RCF for 5 minutes and re-suspended at a density of 6,000 cells/µl in warm Neurobasal-A (Gibco #10888022) + 1X B27 supplement (Gibco #17504044) + 1X GlutaMax (Gibco #35050061) + 1% HEPES (Gibco #15630080) + 1% Penicillin/Streptinomycin (Gibco # 15140122) + 1% fetal bovine serum (Gibco #26140079) + 5 ug/mL Laminin (Sigma-Aldrich #L2020). 150,000 cells were plated on a pre-coated 48-well MEA plate in a 25 uL drop. The day after plating (DIV1), 100% of the media was removed and replaced with warm Neurobasal-A + 1X B27 supplement + 1X GlutaMax + 1% HEPES + 1% Penicillin/Streptinomycin (NBA/B27 medium). Glial growth was not chemically suppressed. Cultures were maintained at 37°C in 5% CO_2_. Media was 50% changed every other day with fresh warm NBA/B27 starting on DIV3, after each recording session.

#### Data analysis of spontaneous recordings

MEA recordings were conducted on media change days prior to media change starting on DIV5. Plates were equilibrated for 5 minutes then recorded for 15 minutes per day using Axion Biosystems Maestro 768 channel amplifier at 37°C in a CO2 gas controlled chamber and Axion Integrated Studios (AxIS) software v2.4. Each well on a 48-well plate is comprised of 16 electrodes on a 4 by 4 grid with each electrode capturing activity of nearby neurons. A Butterworth band-pass filter (200-3000HZ) and an adaptive threshold spike detector set at 7x the standard deviation of the noise was used during recordings. Raw data and a spike list files were collected. Spike list files were used to extract additional spike, burst, and network features, using a custom MEA analysis software package for interpretation of neuronal activity patterns, meaRtools, based on rigorous permutation statistics that enables enhanced identification of over 70 activity features (Gelfman et al., 2018). Specifically, we analyzed spiking and bursting rates, spike density in bursts, periodicity of bursting, burst duration, and the time between bursts (i.e. inter-burst interval, IBI), as well as synchronicity of the network. We determined the parameters for detecting neuronal bursts and network events based on published reports and experimentation (Mack et al., 2014; McConnell et al., 2012). Activity data was inspected to remove inactive electrodes and wells. In order for an electrode to be considered active, we required that at least five spikes per minute were recorded. Wells in which fewer than 4 electrodes were active for > 50% of the days of recording were considered inactive and removed from analyses. For synchronous network events, at least 5 electrodes (> 25% of the total in a well) were required to participate in a network event in order for the network event to qualify as a network spike or burst. Events with less participating electrodes were filtered. Bursts were detected using the Maximum Interval burst detection algorithm (Neuroexplorer software, Nex Technologies) implemented in the meaRtools package (Gelfman et al., 2018). We required that a burst consists of at least 5 spikes and lasts at least 0.05 second, and that the maximum duration between two spikes within a burst be 0.1 second at the beginning of a burst and 0.25 second at the end of a burst. Adjacent bursts were further merged if the duration between them is less than 0.8 second. These parameters were chosen based on the literature and on in-house experimentation (Mack et al., 2014). To analyze data over time, we performed permuted Mann-Whitney U tests. The values for each well for the chosen DIVs were combined and a Mann-Whitney U (MWU) test was performed. The labels for each well (WT vs *Gnb1*^K78R/+^) were then shuffled and permuted 1,000 times to create 1,000 data sets that were tested for significance using a MWU test. We report the permuted p-values as the rank of the true p-value within the distribution of permuted p-values. When mentioned, the *Gnb1*^K78R/+^ well data of each plate was normalized to the WT wells, before combining the plates for statistical analysis using an R script developed in-house. To normalize the data for each plate, we computed the average value amongst WT wells per DIV and then divided the values of each well per DIV by this resulting mean. For comparison, we also combined the individual p-values of each plate using an R script that performs Fisher’s method.

#### Pharmacology

Pharmacological studies were performed on mature cultures between DIV20 and DIV45. Media volumes were equilibrated to 300 µl per well. MEA plates were dosed by addition of a defined amount of a stock concentration of drug to a parallel dosing plate, then 150 uL of the media from the MEA plate was removed and added to the parallel dosing plate and mixed prior to adding the media back to the MEA plate. Spontaneous activity in physiological solution was recorded for 3 min after equilibrating the MEA plates on the Maestro system for 10 min (baseline condition). Then, networks were exposed to the drug and spontaneous activity was recorded 10 min after drug application, for 3 min (drug condition). Results are presented as a ratio drug condition/baseline condition (baseline condition = 1, blue dashed line on graphs in Figures 3-5). To compare the effects of the drugs between WT and *Gnb1*^K78R/+^ neurons, we performed a MWU test with Bonferroni correction.

For the chronic dosing of ethosuximide from DIV3 to DIV31, ethosuximide was added during medium change every other day. It was diluted in pre-warmed NBA/B27 just before medium change to a final concentration of 250 µM, 750 µM or 2 mM from a 200mM stock solution.

### Drugs

All drugs used were purchased from commercial sources: Ethosuximide (Sigma-Aldrich), Valproate sodium (Sigma-Aldrich), Phenytoin (Selleckchem), Gabazine (Tocris), Muscimol (Tocris), Baclofen (Tocris), Bicuculline (Tocris), CGP 55845 (Tocris), NNC 55-0396 (Tocris), ML218 (Tocris), Tertiapin Q (Tocris), 8-CPT (Tocris). For mouse injections, Ethosuximide and Valproate sodium were resuspended in 0.9% saline, Phenytoin in 2% N,N-dimethylacetamide/10% propylene glycol/30% 2-hydroxypropyl-beta-cyclodextrin in H_2_O, and administered via intra-peritoneal injections in a volume of 10 ml/kg. For MEA, drugs were either resuspended in H_2_O for Ethosuximide, Valproate sodium, Muscimol, Baclofen, NNC 55-0396 and Tertiapin Q, or in DMSO for Phenytoin, Gabazine, Bicuculline, CGP 55845, ML218 and 8-CPT. DMSO concentration on MEA never exceeded 0.1%. Concentrations used are reported in figure legends.

### Behavioral experiments

All behavioral experiments were performed in the Mouse NeuroBehavioral Core facility at Columbia University Irving Medical Center.

#### Neonatal pup development

Pups are tattooed in the paws for identification on P2. Developmental milestones and pup vocalizations are conducted on alternated days to avoid over handling.

##### Developmental Milestones

Developmental milestones in pups are important readouts of developmental delays. We measure body weight, righting reflex (the latency to right oneself from a belly-up position to be on all fours), negative geotaxis (on a wire mesh screen, the latency to turn 90 degrees or 180 degrees from a downward-facing start position), and vertical screen holding (the latency too fall off a vertically positioned wire mesh screen). To begin the test, each pup is gently removed from the nest and placed on a clean piece of bench protector. The cage lid is immediately and gently placed back, to reduce agitation in the nest. All assessment is completed by a deft experimenter within 3 min. At the end of the session, the pup is quickly returned to the nest.

##### Ultrasonic Vocalizations

Neonatal mouse pups emit ultrasonic vocalizations (USV) when separated from the dam. Separation-induced vocalizations are tested on postnatal days P5, P7, P9 and P11. The pup is gently removed from the nest and placed in a plastic container (10 cm x 8 cm x 8.5 cm) the bottom of which is covered with a 0.5 cm layer of fresh bedding. The cage lid is immediately and gently placed back, to avoid agitating the dam and the pups in the nest. The container holding the isolated pup is immediately placed inside a sound-attenuating environmental chamber (Med Associates, St. Albans, VT, USA). At the end of the 3 min recording session, each pup is marked (to avoid repeated handling on the same day), and returned to the nest. The experimenter then repeats the same procedure with the next pup, until the whole litter is tested. Ultrasonic vocalizations are recorded by an Ultrasound Microphone (Avisoft UltraSoundGate condenser microphone CM16, Avisoft Bioacoustics, Germany) sensitive to frequencies of 10-180 kHz. Ultrasonic calls are recorded using the Avisoft Recorder software and analyzed using the Avisoft SASLab Pro software.

#### Adult motor tests

##### Open Field exploratory activity

The open field test is the most commonly used general test for locomotor activity. Each mouse is gently placed in the center of a clear Plexiglas arena (27.31 x 27.31 x 20.32 cm, Med Associates ENV-510) lit with dim light (∼5 lux), and is allowed to ambulate freely for 60 min. Infrared (IR) beams embedded along the X, Y, Z axes of the arena automatically track distance moved, horizontal movement, vertical movement, stereotypies, and time spent in center zone. At the end of the test, the mouse is returned to the home cage and the arena is cleaned with 70% ethanol followed by water, and wiped dry.

##### Rotarod

Motor learning was assessed using a mouse accelerating rotarod (Ugo Basile). Mice were placed on the rotating drum that accelerated from 5 to 40 rpm over 5 min. Mice were tested for three trials a day, for 2 consecutive days. The inter-trial interval was 1 h. Rotarod scores were scored for latency to fall or ride the rod around for all three cohorts.

#### Adult learning and memory tests

##### Water maze acquisition and reversal

Spatial learning and reversal learning were assessed in the Morris water maze using procedures and equipment as previously described (Yang et al., 2012, J. Neuro). The apparatus was a circular pool (120 cm diameter) filled 45 cm deep with tap water rendered opaque with the addition of nontoxic white paint (Crayola). Proximal cues were two stickers taped on the inner surface of the pool, approximately 20 cm above the water surface. Trials were videotaped and scored with Ethovision XT 12 (Noldus). Acquisition training consisted of four trials a day for 5 days. Each training trial began by lowering the mouse into the water close to the pool edge, in a quadrant that was either right of, left of, or opposite to, the target quadrant containing the platform. The start location for each trial was alternated in a semi-random order for each mouse. The hidden platform remained in the same quadrant for all trials during acquisition training for a given mouse, but varied across subject mice. Mice were allowed a maximum of 60 s to reach the platform. A mouse that failed to reach the platform in 60 s was guided to the platform by the experimenter. Mice were left on the platform for 15 s before being removed. After each trial, the subject was placed in a cage lined with absorbent paper towels and allowed to rest under an infrared heating lamp for 60 s. Acquisition training continued for 5 d, or until the WT control group reached criterion. Three hours after the completion of last training on day 5, the platform was removed and mice were tested in a 60 s probe trial. Parameters recorded during training days were latency to reach the platform, total distance traveled, and swim speed. Time spent in each quadrant and number of crossings over the trained platform location and over analogous locations in the other quadrants was used to analyze probe trial performance. Reversal training began 3 d after the completion of acquisition training. In reversal training trials, the hidden platform was moved to the quadrant opposite to its location during acquisition training, for each mouse. Procedures for reversal training and probe trial were the same as in the initial acquisition phase.

##### Fear Conditioning

This is a classic test for conditioned learning. Training and conditioning tests are conducted in two identical chambers (Med Associates, E. Fairfield, VT) that were calibrated to deliver identical footshocks. Each chamber was 30 cm × 24 cm × 21 cm with a clear polycarbonate front wall, two stainless side walls, and a white opaque back wall. The bottom of the chamber consisted of a removable grid floor with a waste pan underneath. When placed in the chamber, the grid floor connected with a circuit board for delivery of scrambled electric shock. Each conditioning chamber is placed inside a sound-attenuating environmental chamber (Med Associates). A camera mounted on the front door of the environmental chamber recorded test sessions which were later scored automatically, using the VideoFreeze software (Med Associates, E. Fairfield, VT). For the training session, each chamber is illuminated with a white house light. An olfactory cue is added by dabbing a drop of imitation almond flavoring solution (1:100 dilution in water) on the metal tray beneath the grid floor. The mouse is placed in the test chamber and allowed to explore freely for 2 min. A pure tone (5kHz, 80 dB) which serves as the conditioned stimulus (CS) is played for 30 s. During the last 2 s of the tone, a footshock (0.5 mA) is delivered as the unconditioned stimulus (US). Each mouse received three CS-US pairings, separated by 90 s intervals. After the last CS-US pairing, the mouse is left in the chamber for another 120 s, during which freezing behavior is scored by the VideoFreeze software. The mouse is then returned to its home cage. Contextual conditioning is tested 24 h later in the same chamber, with the same illumination and olfactory cue present but without footshock. Each mouse is placed in the chamber for 5 min, in the absence of CS and US, during which freezing is scored. The mouse is then returned to its home cage. Cued conditioning is conducted 48 h after training. Contextual cues are altered by covering the grid floor with a smooth white plastic sheet, inserting a piece of black plastic sheet bent to form a vaulted ceiling, using near infrared light instead of white light, and dabbing vanilla instead of banana odor on the floor. The session consisted of a 3 min free exploration period followed by 3 min of the identical CS tone (5kHz, 80dB). Freezing is scored during both 3 min segments. The mouse was then returned to its home cage. The chamber is thoroughly cleaned of odors between sessions, using 70% ethanol and water.

### Electrophysiology

Oocyte defolliculation, incubation and RNA injection were performed as described previously (Rishal et al., 2005; Rubinstein et al., 2009). Oocytes were defolliculated with collagenase (Type 1A, Sigma) in Ca-free ND96 solution (see below) and injected with 50 nl of RNA and incubated for 2-4 days in NDE solution (ND96 solution supplemented with 2.5 mM pyruvate and 50 μg/ml gentamicin) at 20°C prior to testing. The standard ND96 solution contained (in mM): 96 NaCl, 2 KCl, 1 MgCl_2_, 1 CaCl_2_, 5 HEPES, and was titrated with NaOH to pH of 7.6-7.8. CaCl_2_ was omitted in Ca^2+^-free ND96.

Whole-cell GIRK and Cav3.2 currents in oocytes were measured using two-electrode voltage clamp (TEVC) with Geneclamp 500 (Molecular Devices, Sunnyvale, CA, USA), using agarose cushion electrodes filled with 3M KCl, with resistances of 0.1–0.8 MΩ for current electrode and 0.2-1.5 MΩ for voltage electrode. To measure GIRK currents we used ND96 solution or high-K+ solution (HK24), in mM: 24 KCl, 72 NaCl, 1 CaCl_2_, 1 MgCl_2_ and 5 HEPES. pH of all solutions was 7.4-7.6. To measure Cav3.2 currents we used high Ba^2+^ solution (Ba40), in mM: 40 Ba(OH)_2_, 50 NaOH, 2 KOH, 5 HEPES, titrated to pH 7.5 with methanesulfonic acid.

Note that to ensure robust activation by Gβγ, the channels were expressed at low surface density of ∼3-5 channels/µm^2^ by using low doses of GIRK1 and GIRK2 RNAs (0.05 ng RNA/oocyte) (see (Peleg et al., 2002; Yakubovich et al., 2015)), which resulted in I_basal_ of ∼ 2-3 µA and maximal Gβγ-evoked currents of ∼8-12 µA with saturating doses of Gβγ for heteromeric GIRK1/2 (Figure 6A,B). In contrast, to obtain comparable surface levels and Gβγ-evoked currents of homotetrameric GIRK2 or GIRK2-YFP (Kahanovitch et al., 2014), we had to inject 2 ng of GIRK2 RNA/oocyte (Figure 6AF,G) (Rubinstein et al., 2009). Note that I_basal_ for GIRK2 was very low, ∼0.05-0.1 µA.

Ethosuximide was purchased from Sigma-Aldrich and dissolved in double-distilled water to get 1 M stock-solution. During the experimental procedure we used 8 concentrations of ETX solutions: in mM: 0.01, 0.03, 0.1, 0.3, 1, 3, 10 and 30. To prepare the experimental solutions we diluted ETX stock-solution in HK24 or Ba40 to 30 mM and the lower concentrations have been prepared by sequential dilutions.

Effect of ethosuximide on GIRK was measured as follows: GIRK currents were measured at a holding potential of −80 mV while the solutions were changed from one to another. First, current was measured in ND96 solution for 5 seconds. Then, solution was switched to HK24 for 30 seconds to measure the basal current. The next solution was HK24 with 0.01 mM ethosuximide for another 30 seconds, the same step was repeated 7 times for each of the ethosuximide solutions. Lastly, HK24 + 2.5 mM Ba^2+^ solution was used to block all GIRK current and to reveal non-GIRK basal current.

Effect of ETX on Cav3.2 was measured as follows from holding potential of −100 mV using the protocol shown in Figure 5F. The protocol was repeated every 10 seconds and solutions were switched to the next does of ethosuximide after stabilization of the current in the previous solution for at least six subsequent measurements.

### Analysis of dose-response relationships

Dose-response data were fitted to the different models using SigmaPlot 11 or 13 (Systat Software Inc., San Jose, CA, USA). The data were fitted to equations 1-3, representing three standard models:

The standard one-component binding isotherm

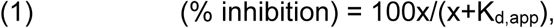

where x is drug concentration, and K_d,app_ is the apparent dissociation constant.

The one-site Hill equation:

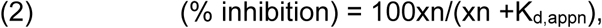

where n is Hill coefficient.

The two-component binding isotherm:

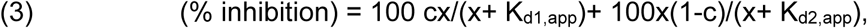

where K_d1,app_ and K_d2,app_ are apparent dissociation constants for 1st and 2nd binding sites, respectively, and c is the fraction of inhibition obtained by the binding of ethosuximide to the high-affinity site.

### Imaging of Gβγ in giant plasma membrane patches

Giant excised PM patches were prepared, stained with antibodies and imaged as described (Singer-Lahat et al., 2000). Oocytes were mechanically devitellinized using fine forceps in a hypertonic solution (in mM: NaCl 6, KCl 150, MgCl_2_ 4, HEPES 10, pH 7.6). The devitellinized oocytes were transferred onto a Thermanox™ coverslip (Nunc, Roskilde, Denmark) immersed in a Ca^2+^-free ND96 solution, with their animal pole facing the coverslip, for 10-20 minutes. The oocytes were then suctioned using a Pasteur pipette, leaving a giant membrane patch attached to the coverslip, with the cytosolic face toward the medium. The coverslip was washed thoroughly with fresh ND96 solution and fixated using 4% formaldehyde for 30 minutes. Fixated giant PM patches were immunostained in 5% milk in phosphate buffer solution (PBS). Non-specific binding was blocked with Donkey IgG 1:200 (Jackson ImmunoResearch, West Grove, PA, USA). Anti-Gβ polyclonal antibody (T-20, from Santa Cruz. #sc-378) was applied at 1:200 dilution, for 45 minutes at 37°C. Anti-rabbit IgG DyLight 649-labeled secondary antibody (1:400; SeraCare Life Sciences, 072-08-18-06) was then applied for 30 minutes in 37°C, washed with PBS, and mounted on a slide for visualization. Imaging was done with Zeiss 510META confocal microscope with a 20x air objective or a 40x water immersion objective using the 633 nm laser in spectral mode, and emission was measured at 673 nm. Imaging of proteins in giant PM patches was performed using the confocal microscope in λ-mode. Images centered on edges of the membrane patches, so that background fluorescence from coverslip could be seen and subtracted.

### Statistical Analyses

Sample size and statistical tests are detailed in the figure legends. Statistical analyses for adult behavioral tests were performed using GraphPad Prism. Two-group comparisons were performed using unpaired two-tailed t-test with correction using the Holm-Sidak method or one-way repeated measures ANOVA with Bonferroni correction. Multiple group comparisons were done using two-way repeated measures ANOVA followed by Sidak’s test. All other statistics were performed in R using a Mann-Whitney U test with 1000 or 10000 permutations, as indicated in legends (See the MEA methods section for more details). For oocyte electrophysiology experiments, statistical analysis was performed on raw data with SigmaPlot 11 or SigmaPlot 13 (Systat Software Inc., San Jose, CA, USA). Two-group comparisons were performed using t-test if the data passed the Shapiro-Wilk normality test and the equal variance test, otherwise a Mann-Whitney Rank Sum Test was used. Multiple group comparisons were done using one-way ANOVA on ranks followed by Dunn’s test.

## SUPPLEMENTAL TABLES

Table S1. MEA plates individual and combined p-values (related to Figure 3)

