## Supplemental Figures for "G protein-coupled potassium channels implicated in mouse and cellular models of GNB1 Encephalopathy"

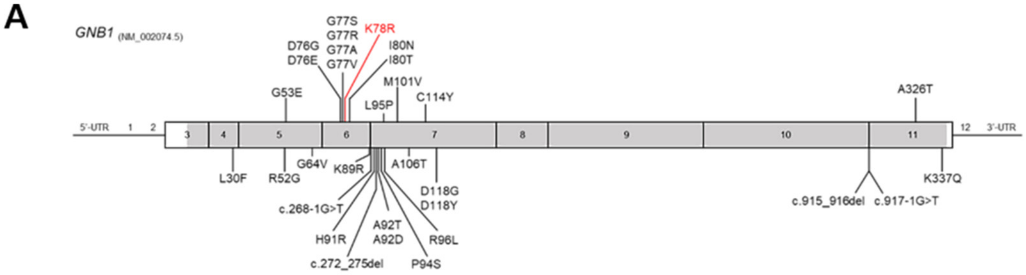


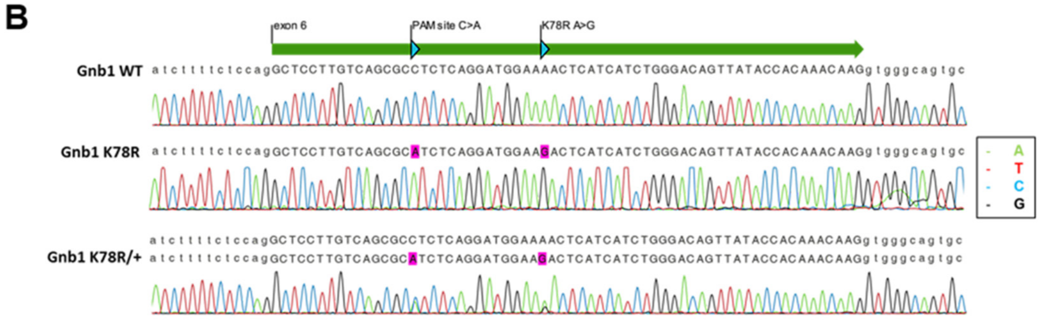


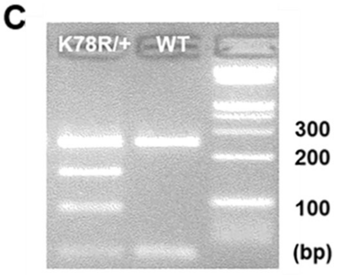


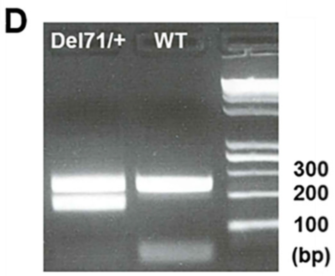


E

**K78R (B6NJ)**

| | WT x Het (P0) | | WT x Het (adults) | | Het x Het (adults) | |
|---|---|---|---|---|---|---|
| | Oberved % (N) | Expected % | Oberved % (N) | Expected % | Oberved % (N) | Expected % |
| WT | 63% (34) | 50% | 79% (129) | 50% | 57% (8) | 25% |
| Het | 37% (20) | 50% | 21% (35) | 50% | 43% (6) | 50% |
| Hom | N/A | N/A | N/A | N/A | 0% (0) | 25% |
| Total | 100% (54) | 100% | 100% (164) | 100% | 100% (14) | 100% |

**K78R (F1H)**

| | WT x Het (adults) | | Het x Het (adults) | |
|---|---|---|---|---|
| | Oberved % (N) | Expected % | Oberved % (N) | Expected % |
| WT | 56% (79) | 50% | 54% (7) | 25% |
| Het | 44% (62) | 50% | 46% (6) | 50% |
| Hom | N/A | N/A | 0% (0) | 25% |
| Total | 100% (141) | 100% | 100% (13) | 100% |

**Del71 (B6NJ)**

| | WT x Het (adults) | | Het x Het (adults) | |
|---|---|---|---|---|
| | Oberved % (N) | Expected % | Oberved % (N) | Expected % |
| WT | 46% (31) | 50% | 37% (10) | 25% |
| Het | 54% (37) | 50% | 63% (17) | 50% |
| Hom | N/A | N/A | 0% (0) | 25% |
| Total | 100% (68) | 100% | 100% (27) | 100% |

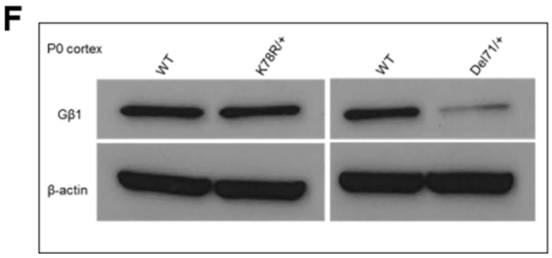


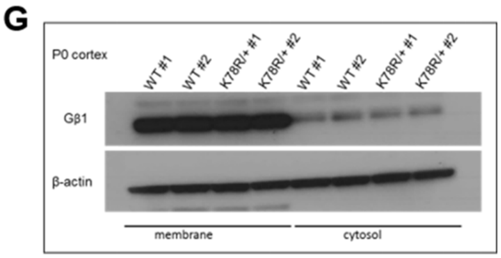

**Figure S1. Generation and characterization of the K78R knock-in mouse model using CRISPR/Cas9 (related to Figure 1)**

**(A)** Schematic of the *GNB1* coding sequence showing the position of published mutations. The K78R mutation located in exon 6 is highlighted in red. **(B)** Sanger sequencing chromatograms showing the WT (top) and K78R (middle) alleles obtained from individual bacterial clones following TOPO TA cloning of the PCR product obtained after amplification of the genomic DNA of a *Gnb1*$^{K78R/+}$ mouse. The third chromatogram corresponds to the direct sequencing of the PCR product, showing the overlay of WT and K78R alleles. **(C)** Typical genotyping result obtained by PCR followed by restriction digest with the HinfI enzyme of tail DNA from a WT and a *Gnb1*$^{K78R/+}$ mouse. **(D)** Typical genotyping result obtained by PCR of tail DNA from a WT and a *Gnb1*$^{Del71/+}$ mouse. **(E)** Numbers and associated percentages of mice of each expected genotypes born from *Gnb1*$^{+/+}$ x *Gnb1*$^{K78R/+}$ or *Gnb1*$^{K78R/+}$ x *Gnb1*$^{K78R/+}$ matings at P0 and after weaning on the B6NJ and F1H backgrounds and from *Gnb1*$^{+/+}$ x *Gnb1*$^{Del71/+}$ or *Gnb1*$^{Del71/+}$ x *Gnb1*$^{Del71/+}$ matings on the B6NJ background. **(F)** Western Blot showing Gβ1 expression in WT, *Gnb1*$^{K78R/+}$ and *Gnb1*$^{Del71/+}$ P0 cortex. β-actin serves as a loading control. Knock-out *Gnb1*$^{Del71/+}$ mice show reduced expression compared to WT mice, as expected, but there is normal Gβ1 expression in *Gnb1*$^{K78R/+}$ mice, suggesting the K78R mutation does not lead to haploinsufficiency. **(G)** Western Blot showing Gβ1 expression in two WT and two *Gnb1*$^{K78R/+}$ P0 cortices following membrane and cytosol fractioning. Most Gβ1 protein is localized in the membrane fraction in both WT and *Gnb1*$^{K78R/+}$ mice.

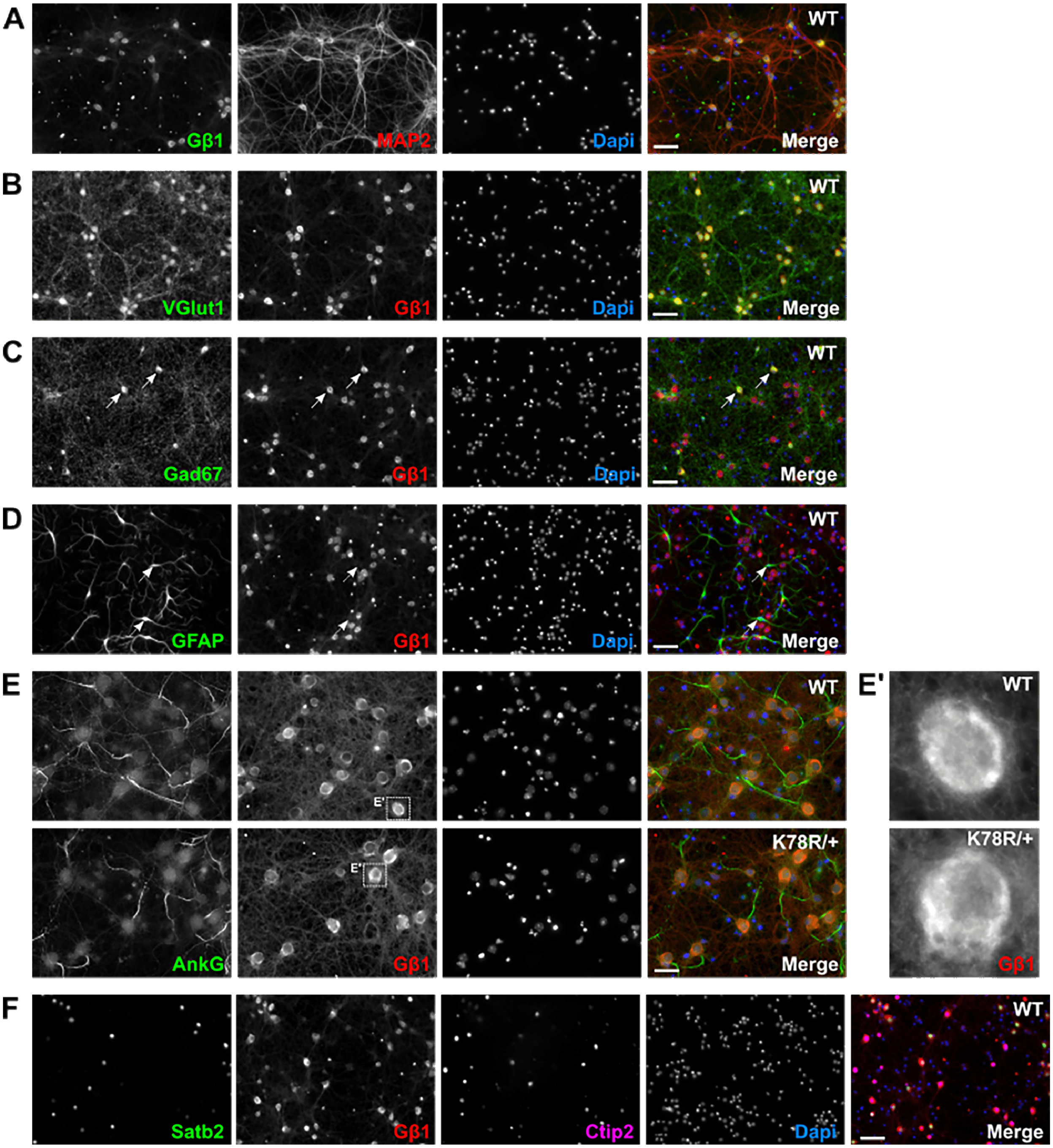


**Figure S2. Gβ1 is strongly localized to the soma of both excitatory and inhibitory neurons (related to Figure 1)** Immunocytochemistry of primary cortical neurons at DIV14. **(A)** Gβ1 (green) is strongly expressed in the soma of all neurons stained by MAP2 (red). **(B)** Gβ1 (red) colocalizes with the excitatory neuron marker, glutamate transporter VGlut1 (green). **(C)** Gβ1 (red) colocalizes with the inhibitory neuron marker, glutamate decarboxylase Gad67 (green). White arrows indicate example of cells showing colocalization. **(D)** Gβ1 (red) does not colocalize with the astrocyte marker GFAP (green). Dashed white arrows indicate example of cells with absence of colocalization. **(E)** Gβ1 (red) is not present in the axon initial segment, visualized with Ankyrin G (green) in both WT and *Gnb1*$^{K78R/+}$ neurons. **(E')** Enlargement of a Gβ1+ neuronal soma from E. **(F)** Gβ1 (red) colocalizes with the deep layer marker Ctip2 (purple) and the upper layer marker Satb2 (green). Scale bars: 50μm for A-D and F; 25μm for E.

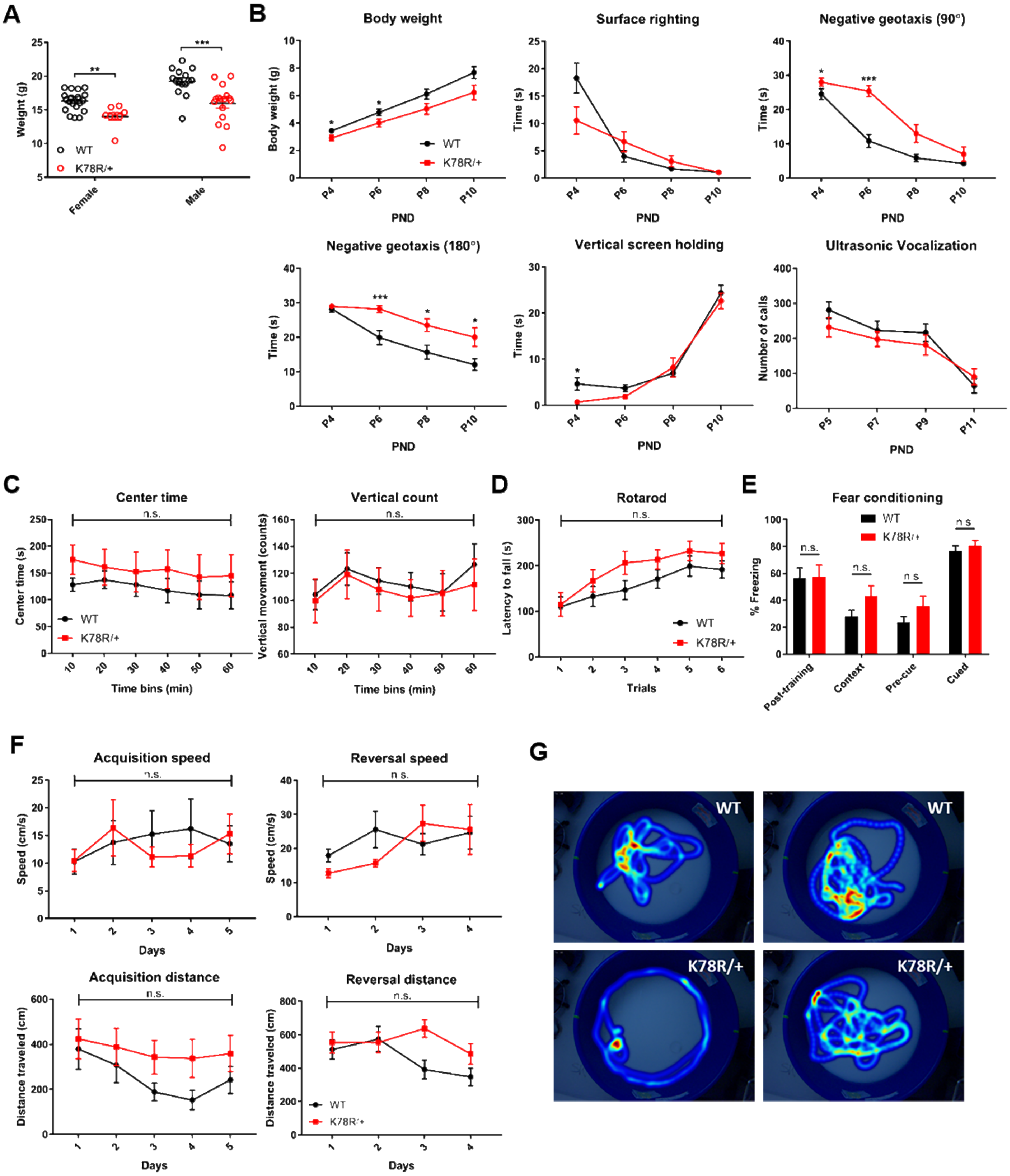

A
Weight (g)
WT
K78R/+
Female
Male
B
Body weight
Body weight (g)
PND
Surface righting
Time (s)
Negative geotaxis (90°)
Negative geotaxis (180°)
Vertical screen holding
Ultrasonic Vocalization
Number of calls
C
Center time
Center time (s)
Time bins (min)
Vertical count
Vertical movement (counts)
n.s.
D
Rotarod
Latency to fall (s)
Trials
E
Fear conditioning
% Freezing
Post-training
Context
Pre-cue
Cued
F
Acquisition speed
Speed (cm/s)
Days
Reversal speed
Acquisition distance
Distance traveled (cm)
Reversal distance
G
WT
K78R/+

**Figure S3. *Gnb1*$^{K78R/+}$ mice present with developmental delay, and motor and cognitive deficits (related to Figure 1)**

**(A)** Weight at 6 weeks old on the B6NJ background. WT (black circles): n = 21 females and 15 males; *Gnb1*$^{K78R/+}$ (red circles): n = 8 females and 17 males; females **p < 0.01 vs. WT and males ***p < 0.001 vs. WT. Mann-Whitney U test with 10000 permutations. **(B)** Developmental milestones between P4 and P10 on the F1H background include body weight, surface righting reflex, negative geotaxis 90º and 180º from downward-facing position, and vertical screen holding: n = 16 WT and 15 *Gnb1*$^{K78R/+}$. Separation-induced USV between P5 and P11: n = 16 WT and 15 *Gnb1*$^{K78R/+}$. Note that we did not observe the inverted U-shaped curve for the WT pups, suggesting that USV may develop earlier on the F1H background. *p < 0.05; ***p < 0.001. Mann-Whitney U test with 10000 permutations**.** **(C)** Additional parameters for open field on the B6NJ background: Vertical exploration and center time. n= 18 WT and 13 *Gnb1*$^{K78R/+}$; n.s.; Two-way repeated measures ANOVA. **(D)** Latency to fall in rotarod test on the B6NJ background. n= 14 WT and 12 *Gnb1*$^{K78R/+}$; n.s.; Two-way repeated measures ANOVA. **(E)** Fear conditioning test for cued and contextual memory. n= 14 WT and 11 *Gnb1*$^{K78R/+}$; n.s.; Unpaired t-test with Holm-Sidak correction for multiple comparisons. **(F)** Control parameters for the Morris Water Maze test on the B6NJ background. Both acquisition and reversal swim speeds and swim distances were unaffected. n = 15 WT and 14 *Gnb1*$^{K78R/+}$; n.s.; Two-way repeated measures ANOVA. **(G)** Additional representative heat maps from the reversal probe trial of the Morris Water Maze test showing a robust difference in search strategy between WT and *Gnb1*$^{K78R/+}$ mice. *All graphs represent mean* ± *SEM*. *n.s., non significant.*

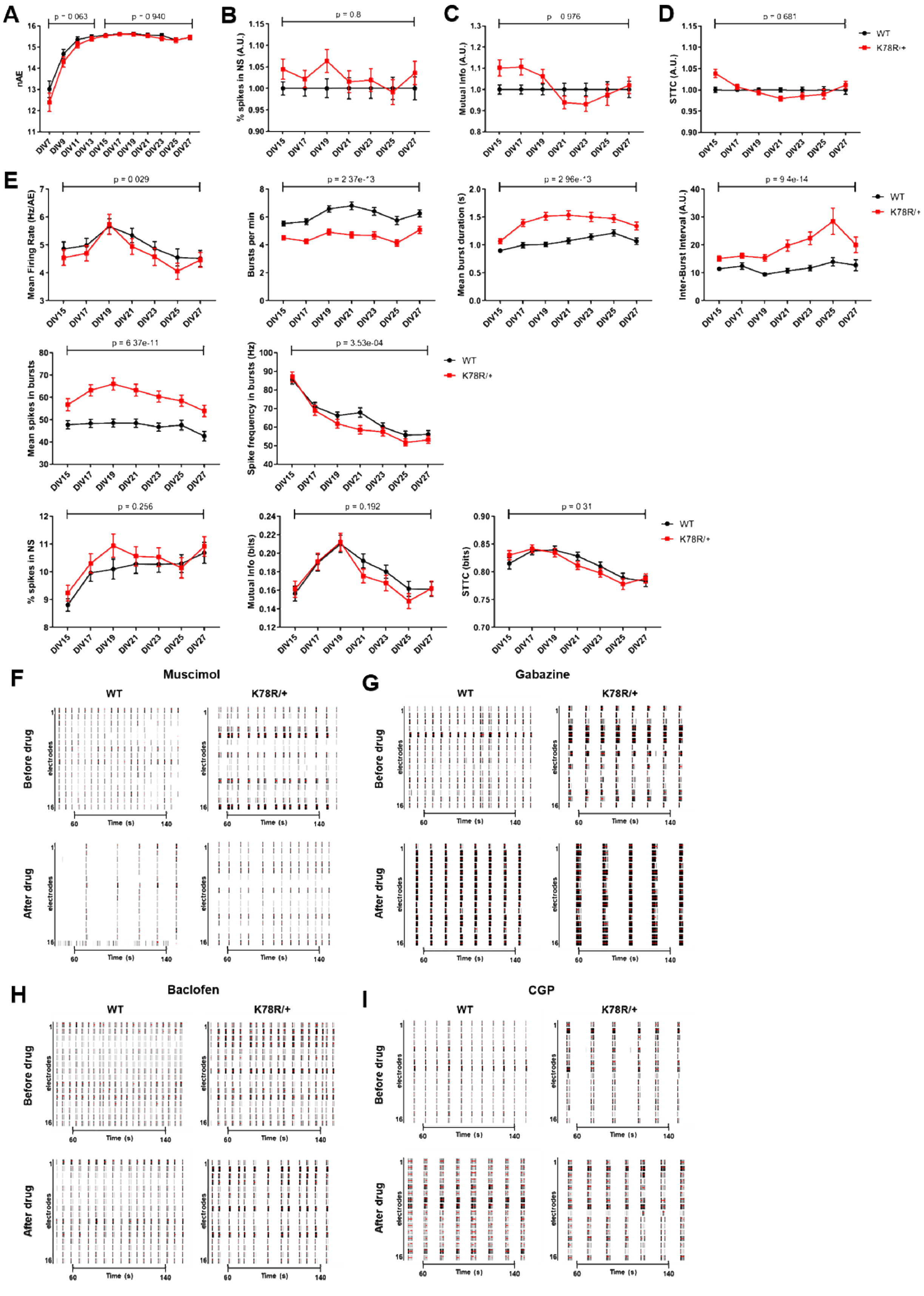

A
B
C
D
E
F
G
H
I
WT
K78R/+
nAE
% spikes in NS (A.U.)
Mutual Info (A.U.)
STTC (A.U.)
Mean Firing Rate (Hz/AE)
Bursts per min
Mean burst duration (s)
Inter-Burst Interval (A.U.)
Mean spikes in bursts
Spike frequency in bursts (Hz)
% spikes in NS
Mutual Info (bits)
STTC (bits)
p = 0.256
p = 0.192
Muscimol
Gabazine
Baclofen
CGP
Before drug
After drug
electrodes
Time (s)

**Figure S4. Excitability phenotypes in *Gnb1*$^{K78R/+}$ cortical neurons (related to Figures 3 and 4)**

**(A)** Graph showing the number of active electrodes (nAE) per well for *Gnb1*$^{K78R/+}$ (red) and WT (black) from DIV7 to DIV27. n = 144 WT wells and 148 *Gnb1*$^{K78R/+}$ wells from 9 plates (8 biological replicates). The combined MWU permutated p-values of each individual plate using Fisher's method is shown. **(B-D)** Graphs representing spontaneous synchrony features on MEA of *Gnb1*$^{K78R/+}$ (red) normalized to WT (black) cortical neurons from DIV15 to DIV27. n = 144 WT wells and 148 *Gnb1*$^{K78R/+}$ wells from 9 plates (8 biological replicates). Permutated p-values were calculated with a Mann-Whitney U test followed by 1000 permutations. **(B)** Percentage of spikes in network spikes (NS). **(C)** Mutual info. **(D)** STTC. **(E)** Graphs showing the combined analysis of the 9 plates from DIV15 to DIV27, as presented in Figure 3, but without normalization to WT. The combined MWU permutated p-values of each individual plate generated using Fisher's method is shown. **(F-I)** Raster plots showing WT and *Gnb1*$^{K78R/+}$ firing before and 10 min after acute drug application across the 16 electrodes of a well of a representative plate, between 50 and 150 seconds of a 3 min recording. The black bars indicate spikes, and the red bars indicate bursts. **(F)** 500nM Muscimol at DIV45. **(G)** 1 µM Gabazine at DIV34. **(H)** 500 nM Baclofen at DIV36. **(I)** 1 µM CGP at DIV36. *All graphs represent mean ± SEM.*

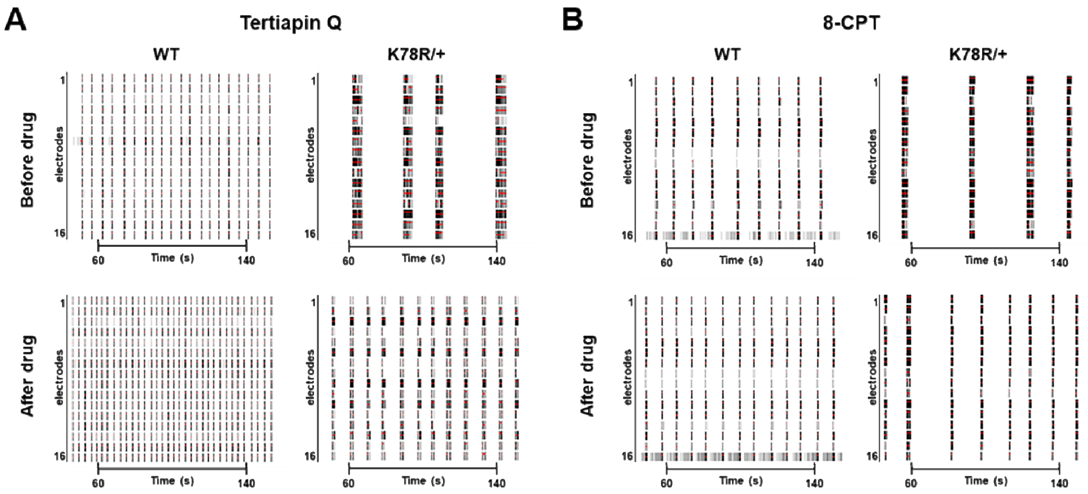


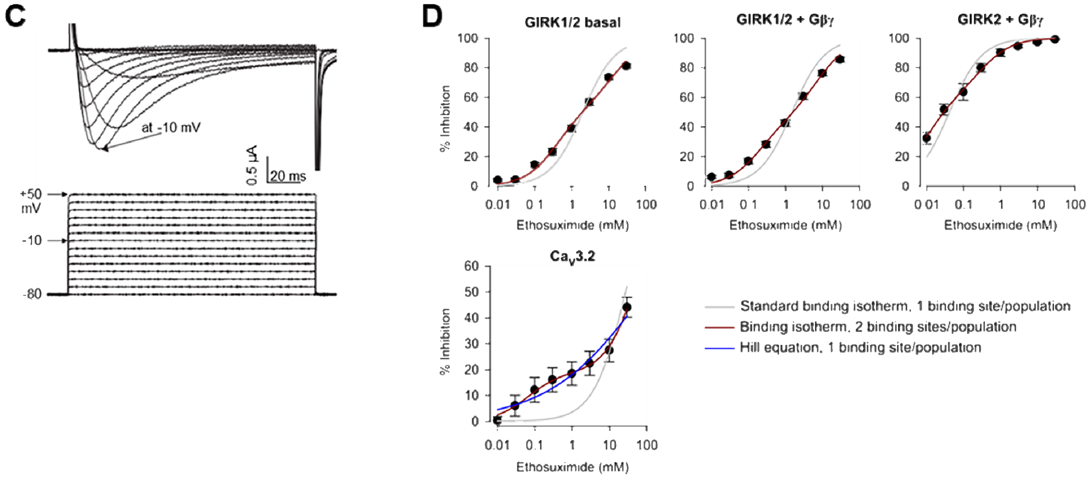


**E**

| Channel | $K_{d,app}$ site 1 mM | Hill coefficient, site 1 | $K_{d,app}$ site 2 mM | Hill coefficient, site 2 | fraction of high-affinity inhibition |
|---|---|---|---|---|---|
| GIRK2 + Gβγ | 0.033 | 0.61 | - | - | |
| GIRK1/2 basal | 2.07 | 0.61 | - | - | |
| GIRK1/2 + Gβγ | 1.5 | 0.6 | - | - | |
| $Ca_v3.2$ | 0.067 | (1) | 69 | (1) | 0.19 |

**F**

| Fits for GIRKs | One-site model | Two-site model | | |
|---|---|---|---|---|
| | $K_{d,app}$ (mM) 1-site model | $K_{d,app}$ site 1 (mM) | $K_{d,app}$ site 2 (mM) | fraction of high-affinity inhibition |
| GIRK1/2 basal | 2 | 0.38 | 12.9 | 0.53 |
| GIRK1/2 + Gβγ | 1.52 | 0.2 | 7.6 | 0.45 |
| GIRK2 + Gβγ | 0.042 | 0.009 | 0.31 | 0.61 |
| Fits for $Ca_v3.2$ | One-site model $K_{d,app}$ (mM) | Hill: $K_{d,app}$ (mM) | Hill coefficient | |
| | 27.7 | 94 | 0.335 | |

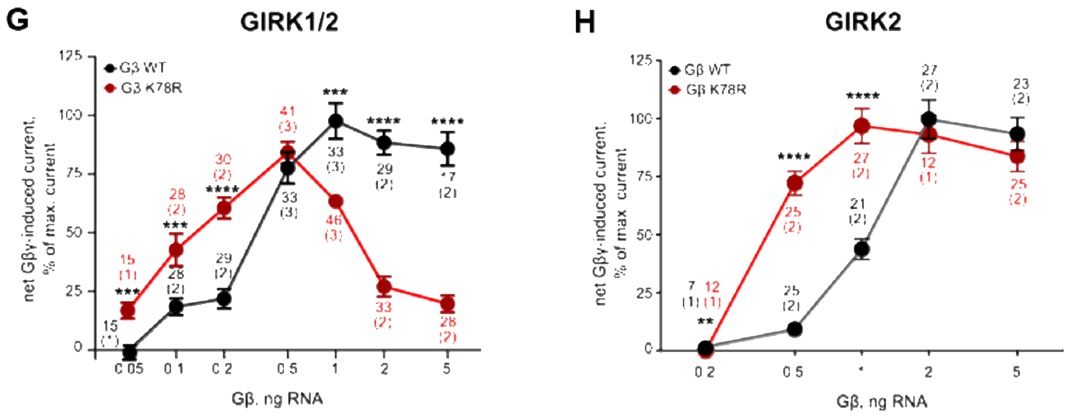

**Figure S5. Electrophysiology in cortical networks on MEA and in *Xenopus* oocytes (related to Figures 5 and 6)**
**(A,B)** Raster plots showing WT and *Gnb1*$^{K78R/+}$ firing before and 10 min after acute drug application across the 16 electrodes of a well of a representative plate, between 50 and 150 seconds of a 3 min recording. The black bars indicate spikes, and the red bars indicate bursts. **(A)** 10 µM Tertiapin Q at DIV33. **(B)** 200 nM 8-CPT at DIV22. **(C)** Representative current-voltage (I-V) relationship of T-type $Ca_V3.2$ channel. Voltage traces are shown on the bottom, current traces on the top. $Ca_V3.2$ yielded typical $I_{Ba}$ with, with a U-shaped current-voltage (I-V) curve (not shown) and maximal current at -10 mV. **(D)** Alternative fitting procedures for dose-response dependencies of ETX inhibition. Data from Figure 5G were fitted to the indicated equations (solid lines). Note that standard one-site binding isotherm did not produce satisfactory fit to any of the datasets. For $Ca_V3.2$, the fit to two-site binding isotherm (shown in Figure 5G) is presented for comparison, to demonstrate that it is superior to both one-site models (standard and Hill). The higher-affinity component had an apparent $K_d$ ($K_{d,app}$) of 67 µM. However, only less than 20% of current was inhibited through this putative high-affinity site. An estimate of 69 mM for $K_{d,app}$ of the low-affinity component was less accurate, because the highest concentration used was 30 mM. **(E)** Parameters of best fits shown in Figure 5D-G. **(F)** Parameters of fitting procedures shown in Figure S5D. **(G,H)** Summary of experiments studying the dose dependence of GIRK activation by $G\beta_{WT}\gamma$ and $G\beta_{K78R}\gamma$. **(G)** Dose-dependent activation of GIRK1/2 by $G\beta_{WT}\gamma$ and $G\beta_{K78R}\gamma$. Net Gβγ-induced currents were calculated in each cell by subtracting $I_{basal}$ of the control group (channel expressed without Gβγ) of the same experiment. To reduce batch-to-batch variability, currents were normalized to the average Gβγ-induced current of the group injected with the highest RNA dose of $G\beta_{WT}\gamma$ in this experiment (2 or 5 ng). Numbers near each experimental point (e.g. 28 (2)) stand for number of oocytes (number of experiments). For each Gβγ RNA dose, currents of the $G\beta_{WT}\gamma$ and $G\beta_{K78R}\gamma$ groups were compared using two-tailed t-test. **$p < 0.01$; ***$p < 0.001$; ****$p < 0.0001$. **(H)** Dose-dependent activation of GIRK2 by $G\beta_{WT}\gamma$ and $G\beta_{K78R}\gamma$, presented as in G.
